# TriTower-m6Am: a triple-tower heterogeneous deep learning architecture integrating semantic, sequential, and structural information for N^6^,2’-O-dimethyladenosine site prediction

**DOI:** 10.64898/2026.08.13.744628

**Authors:** Kaifeng Xiong, Jianhua Jia

## Abstract

**Background:** N^6^,2’-O-dimethyladenosine (m6Am) is a cap-proximal mRNA modification deposited by PCIF1 at the first transcribed nucleotide of eukaryotic mRNAs. Knowing where m6Am sites sit across the transcriptome would help explain how cells tune mRNA stability and translation, but current computational predictors typically depend on a single sequence representation and do not jointly model the semantic, sequential, and structural signals carried by an RNA sequence.

**Results:** We present TriTower-m6Am, a triple-tower architecture that combines three representations: semantic (RNA-FM with BellPooling), sequential (One-Hot BiLSTM), and structural (RGCN with three typed edges). On an independent test set of 640 sequences, TriTower-m6Am reaches AUC = 0.776, MCC = 0.440, and SN = 0.888, against DTC-m6Am’s AUC = 0.765, MCC = 0.411, and SN = 0.800. The 8.8 percentage-point gain in sensitivity means that, for every 100 real m6Am sites, the model recovers roughly 9 additional sites missed by the previous best method. Among the three towers, RGCN alone gives the strongest single signal, and the AUC-weighted ensemble raises sensitivity from the 0.55–0.76 band of the standalone towers to 0.89. Ablating the RGCN edge types shows that backbone connectivity accounts for most of the structural signal.

**Conclusions:** Combining semantic, sequential, and structural views of the same RNA sequence improves m6Am prediction beyond what any single representation achieves. Because each tower’s contribution to the final prediction is a readable voting weight rather than a hidden parameter, the model is not a black box: a user can read off which tower drove a given prediction and trace it back to the corresponding representation, without running a separate post-hoc explainer. The same design pattern can be transferred to other RNA modification site prediction tasks.

**Author summary:** Predicting where m6Am modifications occur on messenger RNA is important for understanding how cells regulate transcript stability and translation. Existing computational methods typically encode the RNA sequence in a single way, such as k-mer counts or a one-hot code, and treat the model as a black box that emits a prediction without explaining which features drove it. We built TriTower-m6Am to address both limitations. Our model combines three independent encoders—a pretrained RNA language model for semantic patterns, a bidirectional LSTM for local nucleotide order, and a relational graph convolutional network for the structural fold—and fuses their outputs by AUC-weighted voting, so the contribution of each tower to a given prediction is a readable number rather than a hidden parameter. On an independent benchmark the ensemble improves sensitivity by 8.8 percentage points over the prior best method, and ablating the graph’s edge types reveals that linear backbone connectivity, rather than long-range base-pairing, carries most of the structural signal. The same triple-tower pattern can be transferred to other RNA modification site prediction tasks.

## Introduction

Of the chemical modifications on eukaryotic mRNA, m6Am (N^6^,2’-O-dimethyladenosine) is unusual in that it is found only at the first transcribed nucleotide, and only when that nucleotide is an adenosine [1, 2]. The modification is written by PCIF1 (also called CAPAM) and can be erased by FTO [3, 4], which places m6Am among the reversible epitranscriptomic marks. Its cap-proximal position gives it a disproportionate functional role: m6Am reduces decapping, lengthens transcript half-life, and modulates translation, splicing, and disease-associated phenotypes [5–8]. This is distinct from internal marks such as m6A or m5C, which decorate the transcript body without the same positional constraint.

Mapping m6Am at single-nucleotide resolution relies on antibody-based assays, chiefly miCLIP and m6Am-seq [3, 9]. These experiments are expensive and low-throughput, which has motivated a series of computational predictors. DTC-m6Am frames the task as an imbalanced-classification problem and pairs DenseNet with an attention module [10]. Deep-m6Am feeds pseudo nucleotide composition into a deep network and uses SHAP to prune features [11]. im6Am-DC swaps in coordinate attention and focal loss to cope with the skewed class ratio [12]. Each of these tools works, but each also represents the input sequence in only one way: k-mer statistics, hand-crafted physicochemical features, or a one-hot code. Three information channels that an RNA sequence carries—the semantic patterns learned by large-scale pre-training, the local sequential order of nucleotides, and the structural fold encoded by base-pairing and neighborhood relationships—are not used together in any of them. Secondary structure, in particular, is largely set aside, even though the local fold governs how PCIF1 reaches its substrate. The third shortcoming is opacity: a prediction is emitted, but the feature basis for that prediction is not.

Pre-trained RNA language models such as RNA-FM [13] and RNAErnie [14] offer one way to recover the semantic channel: trained on millions of sequences, they encode co-occurrence patterns that k-mer counts miss. They have not, however, been combined with an explicit sequential encoder and a structural graph in a single m6Am predictor, and the question of whether the three channels are complementary or redundant at this task has therefore not been answered.

**TriTower-m6Am** is built around that question. The model has three independent towers. The RNA-FM tower reads a 41-nt window with a pre-trained language model and pools the per-position embeddings with a learnable bell-shaped attention (BellPooling) centered on the candidate site. The One-Hot BiLSTM tower consumes the same window as a 4-dimensional one-hot code and processes it bidirectionally, keeping the hidden state at the central position as its sequential summary. The RGCN tower builds a graph over the 41 nucleotides with three edge types—backbone, base-pairing, and neighborhood—and runs a relational graph convolution that assigns each edge type its own weight matrix. The three tower outputs are fused by AUC-weighted voting, so the contribution of each tower is a readable number rather than a hidden parameter. On the DTC-m6Am benchmark (640 balanced independent test sequences), the ensemble improves on the prior best method most visibly in sensitivity. Because the RGCN distinguishes edge types, the same ablation that validates the design also exposes which structural relationships the model relies on, so interpretation is a by-product of the architecture rather than an additional stage. In summary, TriTower-m6Am addresses the three shortcomings identified above with three explicit strategies: (1) a pretrained RNA language model (RNA-FM) to capture the semantic patterns that k-mer counts miss; (2) a BiLSTM to encode the local nucleotide order that one-hot convolutions treat only implicitly; and (3) a relational graph convolutional network with typed edges to model the structural fold, while the AUC-weighted voting weights make each tower’s contribution to a prediction directly auditable rather than opaque.

## Materials and methods

### Dataset

All experiments use the DTC-m6Am benchmark released by Huang et al. [10]. A sample is a 41-nt window centered on the candidate site (position 20, 0-indexed), with 20 nt of upstream and downstream context.

Two key constraints shape the dataset. First, the central nucleotide is adenosine in every sample, positive or negative, so the model cannot classify by reading the candidate base itself; it must extract signal from the flanking context. Second, the positives are miCLIP-confirmed m6Am sites, while the negatives are adenosine positions that tested negative in the same miCLIP experiments, so the negative label is experimentally verified rather than drawn from arbitrary adenosine sites.

The training set holds 40,700 windows (3,700 positives, 37,000 negatives; *≈*10:1 ratio), and the held-out test set contains 640 windows balanced at 320 each (Table 1). The skewed ratio is biological in origin: among cap-proximal adenosines, only a minority carry the modification. Secondary structure for each window was predicted with RNAfold from the ViennaRNA package [15] and stored as dot-bracket strings, which the RGCN tower later parses into pairing and loop annotations.

**Table 1.** Dataset split and class distribution. “Pos.” = positive samples; “Neg.” = negative samples. The test set is balanced to give an unbiased estimate of per-class metrics.

| Split | Pos. | Neg. | Total | Pos:Neg |
| --- | --- | --- | --- | --- |
| Training | 3,700 | 37,000 | 40,700 | $\approx 1:10$ |
| Test | 320 | 320 | 640 | 1:1 |

### Overall architecture

The model is organized as three independent towers whose predictions are fused at the end (Fig 1). As sketched in Fig 1a, the pipeline runs from raw 41-nt window through representation extraction, 5-fold cross-validated training of each tower, and finally AUC-weighted voting.

**Fig 1.**
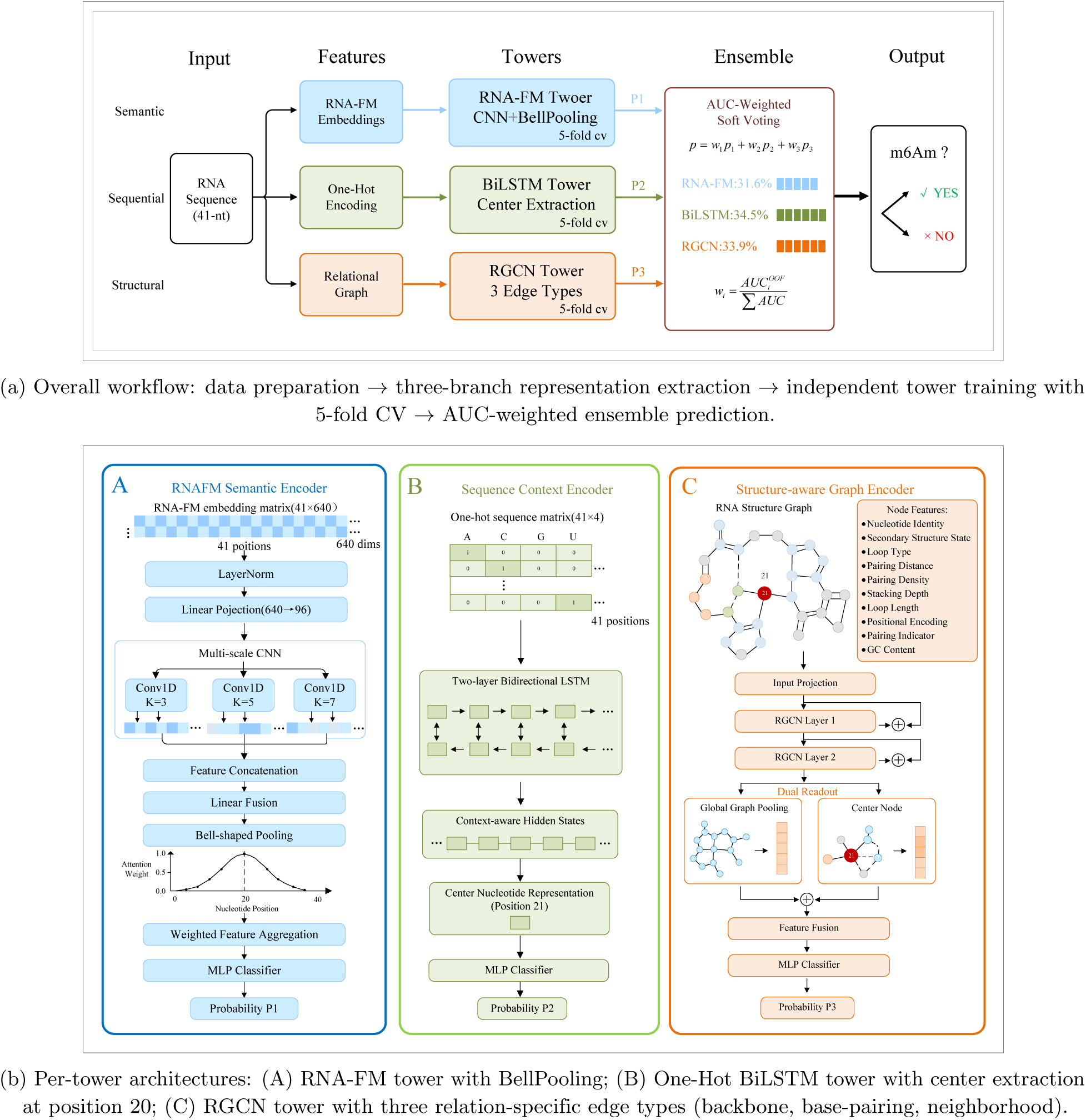
Overview of the TriTower-m6Am framework. (a) End-to-end workflow from input sequence to ensemble prediction. (b) Architecture of the three independently trained towers, each operating on a distinct representation of the 41-nt input.

Fig 1b shows the per-tower encoders. Each tower consumes the same input window but through a different representation, so the three towers answer different questions about the same sequence: the RNA-FM tower asks what semantic patterns the window carries, the One-Hot BiLSTM tower asks how the nucleotide order reads, and the RGCN tower asks how the nucleotides are arranged in 2D space.

Keeping the towers separate during training has three practical benefits: the optimization landscape of each tower stays small, each tower can be ablated cleanly, and each tower is free to specialize in its own representation space without being pulled toward a shared objective.

### RNA-FM semantic tower

The first tower is built on RNA-FM [13], a Transformer-based [16] RNA language model pre-trained on 23.7 million ncRNA sequences. For every 41-nt window, we pre-compute the per-nucleotide embedding from the frozen RNA-FM backbone, which produces a 41 *×* 640 matrix. The downstream encoder then squeezes this into a 48-dimensional sequence representation through six stages: (1) layer normalization on the raw 640-dim embeddings; (2) a linear projection 640 *→* 96 with GELU; (3) three parallel 1-D convolutions (kernels 3, 5, 7), each yielding 48 channels followed by BatchNorm and GELU, so that local patterns at three scales are read in parallel; (4) concatenation of the three branches (144 channels) and projection back to 48; (5) BellPooling, a position-aware weighted sum that converts the 41 positions into a single vector; and (6) a two-layer classifier Linear(48 *→* 32) *→* GELU *→* Linear(32 *→* 1).

BellPooling is what makes this tower position-aware. The 41 per-position features are summed with learned weights *w_i_* produced by

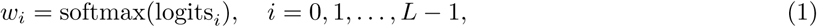

with *L* = 41. Before training, the logits are filled from a bell-shaped Cauchy-type prior

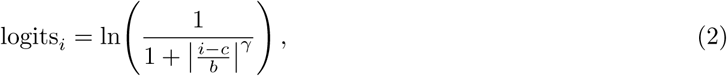

centered at *c* = 20 with width *b* = 5 and exponent *γ* = 2. Under softmax, this gives *w_i_ ∝* 1*/*(1 + *|*(*i −* 20)*/*5*|*^2^), i.e. a symmetric bell that peaks at the candidate site and is free to shift during training. The prior encodes a simple expectation (positions next to the cap-proximal adenosine are more informative) which training can then refine.

### One-Hot BiLSTM sequential tower

The second tower drops every external prior and reads the window as a raw 4-dimensional one-hot code (A = [1, 0, 0, 0], U = [0, 1, 0, 0], G = [0, 0, 1, 0], C = [0, 0, 0, 1]), giving a 41 *×* 4 input. The encoder is a 2-layer bidirectional LSTM [17] with 64 units per direction (128 total), so each position’s hidden state is informed by nucleotides on both sides. As the representative vector we keep only the hidden state at position 20: the candidate site sits in the middle of the window, and the bidirectional state at that position already aggregates upstream and downstream context. A two-layer classifier Linear(128 *→* 64) *→* GELU *→* Linear(64 *→* 1) then maps this 128-dim summary to a logit. The design is deliberately low-information on the input side—no pre-training, no structure—so whatever signal this tower recovers must come from nucleotide order itself, which is what we want to measure against the other two towers.

### RGCN structural tower

The third tower turns the secondary structure into a graph and processes it with a Relational Graph Convolutional Network [18]. We chose RGCN over the more common GAT [19] for one practical reason: GAT shares one attention mechanism across every edge type, so it cannot tell us whether the model is reading the backbone, the base pairs, or the local neighborhood. RGCN, by giving each edge type its own weight matrix, keeps these contributions separate, which is exactly what the later edge-type ablation needs.

#### Graph construction

Each RNA sequence is represented as a graph with 41 nodes (one per nucleotide). Three types of directed edges are defined: *backbone edges* (type 0) connect adjacent nucleotides (*i, i* + 1), capturing sequential connectivity (80 directed edges per sequence); *base-pairing edges* (type 1) connect nucleotides paired in the predicted secondary structure, following the standard RNA pairing rules including the Watson–Crick pairs A–U and G–C and the thermodynamically stable wobble pair G–U that frequently occurs in RNA hairpins and stems; and *neighborhood edges* (type 2) connect nucleotides at distance 2 (*i, i* + 2), capturing near-neighbor interactions (78 directed edges per sequence).

#### Node features

Each node is represented by a 21-dimensional structural feature vector (Table 2).

**Table 2.** Structural node features (21 dimensions).

| Feature group | Dim | Description |
| --- | --- | --- |
| Nucleotide one-hot | 4 | A, U, G, C |
| Secondary structure | 3 | Paired (, paired ), unpaired . |
| Loop type | 6 | Hairpin, interior, bulge, multi, external, stem |
| Pair distance | 1 | Distance to paired partner (0 if unpaired) |
| Pair density | 1 | Number of paired neighbors within window |
| Stacking depth | 1 | Normalized consecutive base-pair stacking depth |
| Loop length | 1 | Normalized length of containing unpaired region |
| Position (sin, cos) | 2 | Sinusoidal positional encoding |
| Is paired | 1 | Binary indicator |
| GC content | 1 | Local GC content |
| **Total** | **21** |  |

### RGCN architecture

The tower consists of four stages: (1) input projection: Linear(21 *→* 64), followed by LayerNorm, GELU activation, and Dropout(0.3); (2) two consecutive RGCNConv(64, 64, num relations = 3, num bases = 3) layers, each augmented by residual connections and LayerNorm; (3) readout: concatenation of global mean pooling and the center node embedding, projected to 64 dimensions via a linear layer; and (4) classification: Linear(64 *→* 32), GELU activation, and a final Linear(32 *→* 1). The RGCN uses 3 basis decomposition matrices (num bases = 3) to control parameter complexity. Edge dropout (rate = 0.1) is applied during training for regularization.

### Implementation

TriTower-m6Am is implemented in Python 3.9 with PyTorch 2.0 [20] and PyTorch Geometric (PyG) [21] for RGCN operations. RNA secondary structure is predicted using RNAfold from the ViennaRNA package 2.0 [15]. RNA-FM embeddings are extracted using the pre-trained model from Chen et al. [13]. All experiments run on a single NVIDIA GPU with 16 GB of memory. The source code is available at https://github.com/xiong-0212/TriTower-m6Am.

### Training and ensemble strategy

#### Loss function and class imbalance

We use BCEWithLogitsLoss with a positive class weight min(*n_neg_/n_pos_,* 3.0) to address the *∼*10:1 class imbalance. The cap at 3.0 was chosen empirically to prevent positive samples from dominating the loss; higher caps caused training instability and specificity degradation, while this cap maintained a stable sensitivity–specificity balance across all towers. Label smoothing (*y_smooth_* = *y ×* 0.92 + 0.04) softens the target distribution and improves calibration under class imbalance.

#### Optimization

We use AdamW [22, 23] with learning rate 5 *×* 10*^−^*^4^ and weight decay 5 *×* 10*^−^*^3^. The learning rate follows a linear warmup over the first 10 epochs, then decays via cosine annealing [24]. We clip gradients at a maximum norm of 1.0 and train with automatic mixed precision (AMP) using PyTorch’s GradScaler.

Early stopping uses balanced validation MCC computed on a 1:1 downsampled subset of the validation fold, with patience of 50 epochs. Optimal hyperparameters were identified through a two-phase grid search. Phase 1 screened 144 configurations on a coarse grid. Phase 2 validated the top-5 configurations with full 5-fold cross-validation. The final values are listed in Table 3. The independent test set was never used during hyperparameter search; it was reserved exclusively for final evaluation.

**Table 3.** Optimal hyperparameters identified by two-phase optimization.

| Hyperparameter | Search Space | Optimal |
| --- | --- | --- |
| Learning rate | {1e-4, 3e-4, 5e-4, 1e-3} | 5e-4 |
| Weight decay | {1e-3, 5e-3, 1e-2} | 5e-3 |
| Dropout | {0.2, 0.3, 0.4, 0.5} | 0.4 |
| EMA decay | {0.99, 0.995, 0.999} | 0.990 |
| *Fixed training parameters* |  |  |
| Label smoothing | — | $y \times 0.92 + 0.04$ |
| Positive class weight cap | — | 3.0 |

We apply an exponential moving average (EMA, decay 0.99) of model parameters during training; the smoothed shadow parameters are used for validation and inference.

#### Cross-validation

Each tower is trained with 5-fold stratified cross-validation (random seed = 123). The training set is split into five folds stratified by class label (positive/negative) using sklearn.model selection.StratifiedKFold with shuffle=True and random state=123, which preserves the *∼*10:1 negative-to-positive ratio within each fold. For each fold, the model is trained on 4/5 of the training set (32,560 samples) and validated on the held-out 1/5 (8,140 samples). The out-of-fold (OOF) predictions are collected for all training samples, and the test set predictions from all 5 folds are averaged. The independent test set (640 samples) is held out from this procedure and used only for final evaluation.

#### AUC-weighted ensemble

The three towers are ensembled via AUC-weighted voting. The ensemble weight for tower *i* is

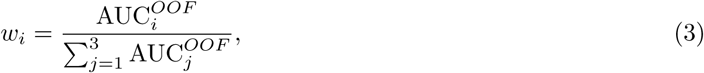

where 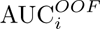 is the OOF AUC of tower *i* on the training set. The final ensemble probability is

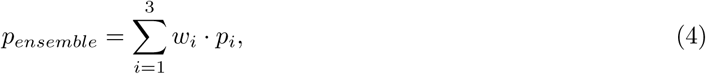

a standard weighted averaging scheme for classifier ensembles [25, 26].

### Evaluation metrics

Performance is evaluated using AUC (Area Under the ROC Curve), MCC (Matthews Correlation Coefficient), SN (Sensitivity/Recall), SP (Specificity), and F1 Score, all computed on the independent test set. The MCC is defined as

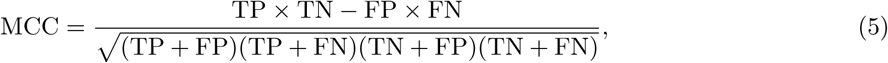

where TP, TN, FP, FN denote true positives, true negatives, false positives, and false negatives, respectively. Threshold-dependent metrics (MCC, SN, SP, F1, ACC) are reported at the optimal threshold determined by maximizing balanced validation MCC on OOF training predictions. The default threshold of 0.5 is used only when comparing against published methods whose threshold-dependent metrics are reported at 0.5.

## Results

### Overall performance and ROC/PR analysis

Table 4 reports the performance of TriTower-m6Am against DTC-m6Am, three published m6Am predictors, and three deep learning baselines. TriTower-m6Am achieves the highest AUC (0.776), MCC (0.440), and F1 (0.750) on the independent test set.

**Table 4.** Performance comparison on the independent test set (640 samples). Best values in bold. *Results from the DTC-m6Am study [10], evaluated on the same independent test set.

| Method | AUC | MCC | SN | SP | F1 |
| --- | --- | --- | --- | --- | --- |
| **TriTower-m6Am (ours)** | **0.776** | **0.440** | **0.888** | 0.522 | **0.750** |
| DTC-m6Am [10] | 0.765 | 0.411 | 0.800 | 0.530 | 0.740 |
| m6AmPred* [27] | 0.735 | 0.289 | 0.887 | 0.358 | 0.702 |
| DLm6Am* [28] | 0.730 | 0.322 | 0.875 | 0.409 | 0.710 |
| m6Aminer* [29] | 0.754 | 0.331 | 0.874 | 0.420 | 0.713 |
| CNN baseline | 0.765 | 0.400 | 0.713 | 0.688 | 0.704 |
| BiLSTM baseline | 0.746 | 0.382 | 0.616 | 0.763 | 0.664 |
| Transformer baseline | 0.724 | 0.360 | 0.709 | 0.650 | 0.689 |

The most telling column is sensitivity. At SN = 0.888, the model recovers 88.8% of true m6Am sites, 8.8 percentage points above DTC-m6Am’s 0.800. Translated to the bench, this means that for every 100 real m6Am sites, our model finds roughly 9 sites that the previous best method would have missed. For a wet-lab pipeline in which each missed site is a separate validation experiment, that is the gap that matters most, and it is wider than the AUC gap (0.011) suggests at first glance.

We re-evaluated DTC-m6Am from its official checkpoint under identical preprocessing and reproduced its reported AUC = 0.765 and MCC = 0.411, so the comparison is against the actual model rather than literature numbers. The sensitivity gain is not bought at the price of specificity: TriTower-m6Am’s SP = 0.522 sits alongside DTC-m6Am’s SP = 0.530, and both are modest—a reminder that, on a balanced test set with 320 positives and 320 negatives, roughly half of the non-m6Am adenosines are still misread.

Three standard deep learning baselines (CNN, BiLSTM, Transformer) trained on the same one-hot input with the same hyperparameters all trail the ensemble. The CNN is the strongest of the three (MCC = 0.400), followed by the BiLSTM (0.382) and the Transformer (0.360). We read this as evidence that the lift does not come from picking the right architecture in isolation—it comes from combining heterogeneous representations, since no single encoder in the comparison clears MCC = 0.40 on its own while the ensemble reaches 0.44.

Fig 2 visualizes the multi-metric comparison. The sensitivity advantage of TriTower-m6Am over all competing methods is immediately apparent in the radar profile. The AUC, MCC, and F1 advantages are also visible, confirming that the multi-representation ensemble captures complementary signals that no single architecture can exploit in isolation. The ensemble processes one sample in 6.18 ms with 329K parameters (S1 Table), well within commodity GPU capacity.

**Fig 2.**
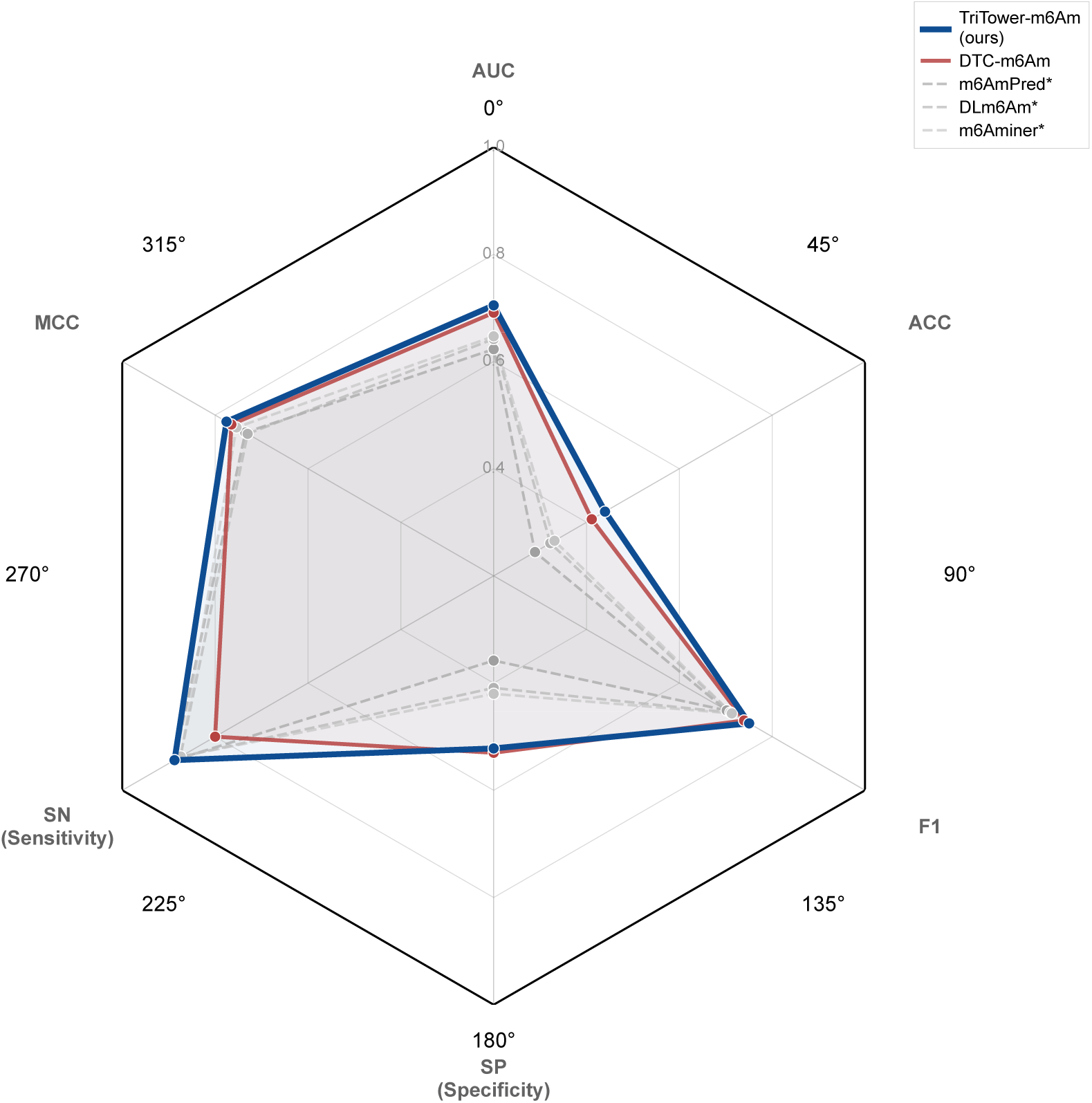
Radar chart comparison of model performance across six metrics. Values closer to the outer circle indicate better performance. *Results from the DTC-m6Am study [10].

The ROC and PR curves (Fig 3) show that the ensemble achieves the highest overall AUC (0.776) and AP (0.758; Average Precision, the area under the PR curve). Among the three towers (Table 5), RGCN gives the strongest standalone AUC (0.768), with RNA-FM close behind (0.762) and BiLSTM last (0.756). What matters for ensembling, though, is not the AUC ranking but the shape of each tower’s error. RNA-FM errs on the side of calling positives (SN high, SP low), BiLSTM errs the other way (SN low, SP high), and RGCN sits between the two. This is the configuration ensemble methods are designed for: the towers make mistakes in different directions, so a disagreement between two towers carries information that the third tower can break, and an agreement between any two is usually a correct call. The ensemble converts this into SN = 0.888 with SP = 0.522—higher sensitivity than any single tower, at a specificity that is still usable for downstream filtering.

**Fig 3.**
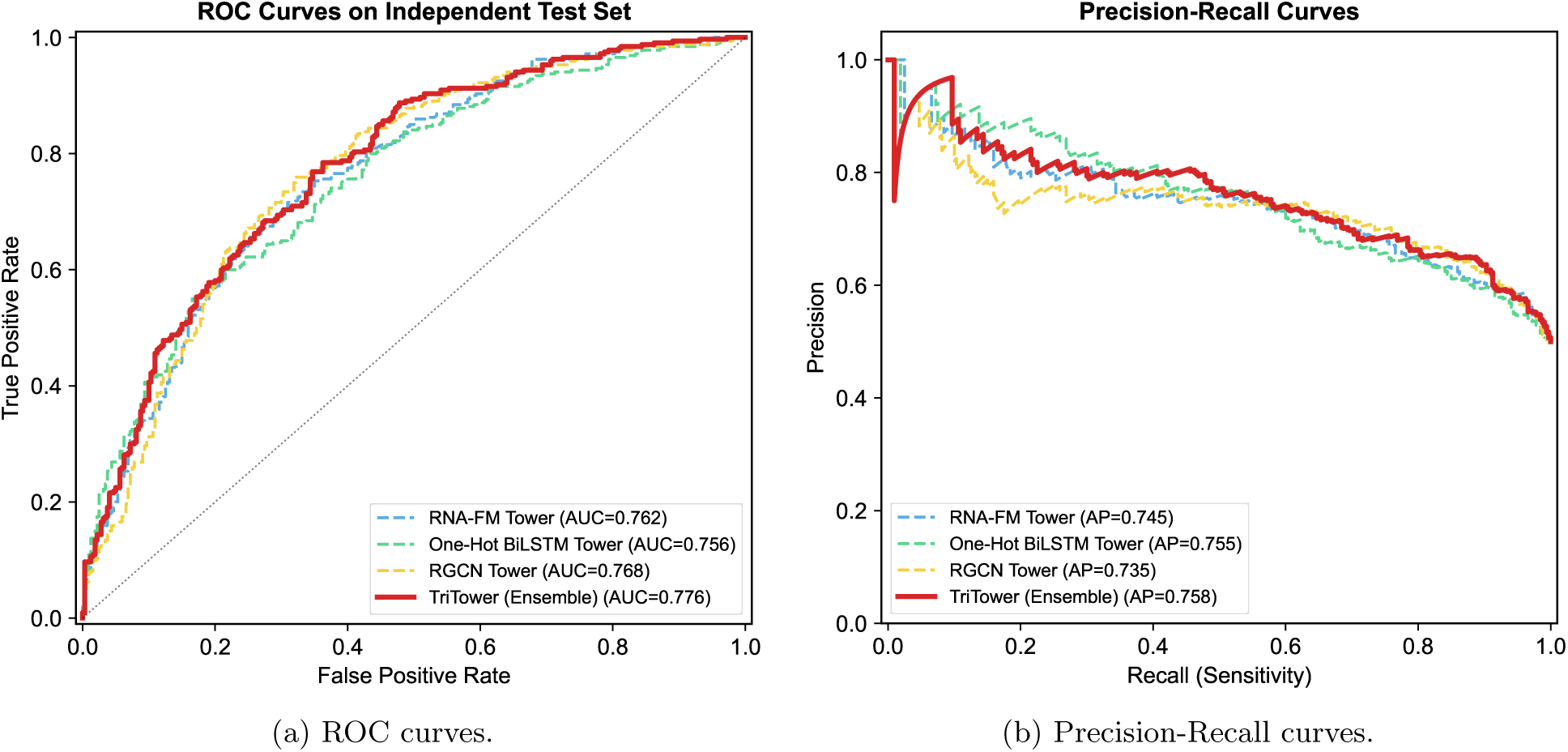
ROC and Precision-Recall curves on the independent test set. Solid lines represent the ensemble (red) and individual towers. AUC and AP values are listed in the legends.

**Table 5.** Tower ablation study. Checkmarks indicate which towers are included. Left columns ablate one tower from the full ensemble; right columns evaluate each tower in isolation.

|  |  |  |  |  |  |  |  |
| --- | --- | --- | --- | --- | --- | --- | --- |
| RNA-FM | ✓ |  | ✓ | ✓ | ✓ |  |  |
| One-Hot BiLSTM | ✓ | ✓ |  | ✓ |  | ✓ |  |
| RGCN | ✓ | ✓ | ✓ |  |  |  | ✓ |
| AUC | **0.776** | 0.770 | 0.773 | 0.769 | 0.762 | 0.756 | 0.768 |
| MCC | **0.440** | 0.426 | 0.434 | 0.398 | 0.399 | 0.401 | 0.439 |
| SN | **0.888** | 0.881 | 0.713 | 0.547 | 0.744 | 0.550 | 0.759 |
| SP | 0.522 | 0.516 | 0.722 | 0.834 | 0.653 | 0.834 | 0.678 |

**Fig 4.**
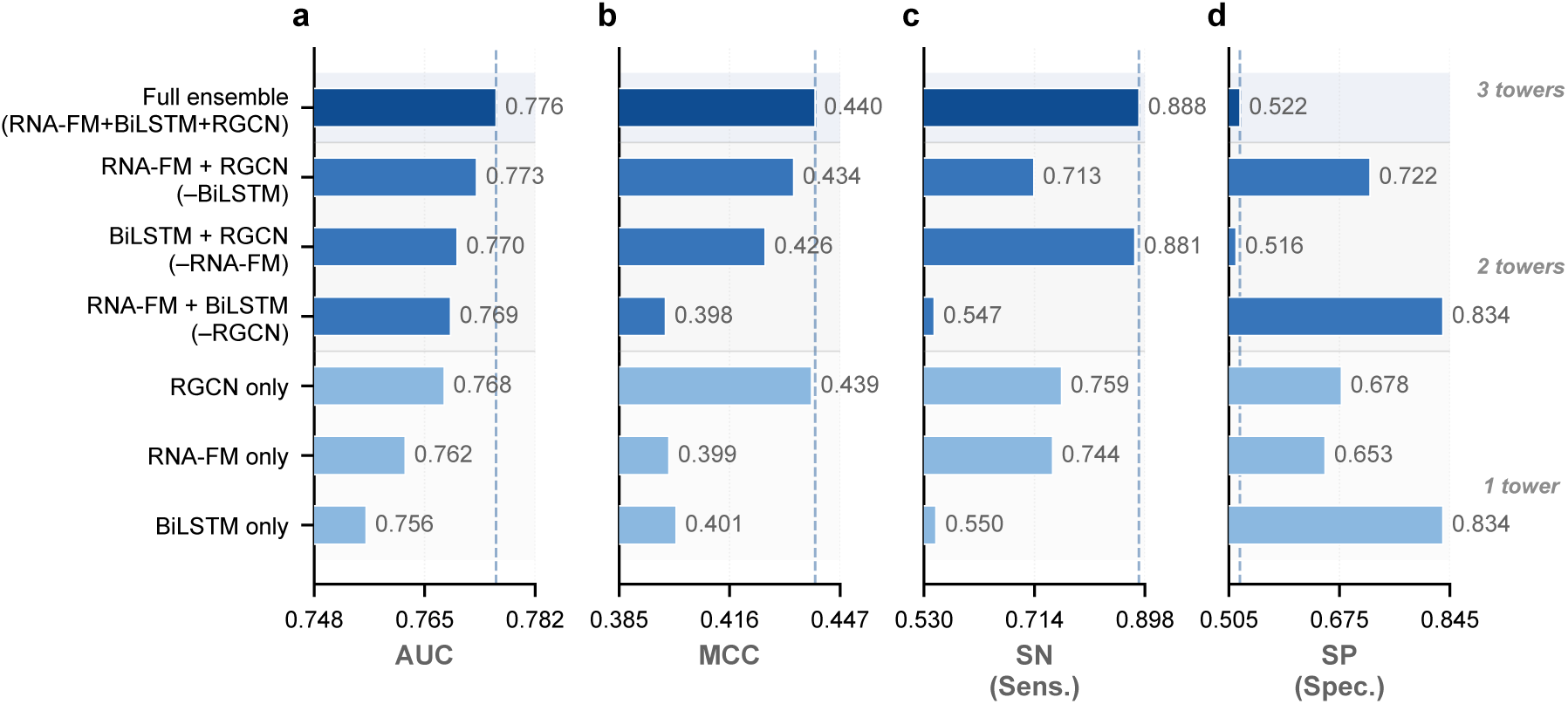
Tower ablation: per-metric comparison. Horizontal bars show AUC, MCC, SN, and SP for each tower configuration. The dashed vertical line marks the full-ensemble value. Configurations are grouped by number of towers (3, 2, 1).

### Representation complementarity: ablation and parameter-matched baselines

Overall performance tells us that the ensemble works; it does not tell us why. The ablations in this section pull the model apart at two levels: first at the tower level (which representation matters?) and then at the component level (which design choices inside each tower matter?).

Table 5 reports the tower ablation. RGCN is the strongest standalone tower (AUC = 0.768, MCC = 0.439) and also the most balanced in its sensitivity–specificity trade-off, which fits the intuition that the local fold around the cap-proximal adenosine is informative for PCIF1 recognition. RNA-FM matches RGCN in AUC (0.762) but tilts toward sensitivity, which we attribute to the pre-trained semantic prior: it ranks cap-proximal contexts the way it has seen them in millions of ncRNAs, and this tends to flag candidate positives aggressively. The One-Hot BiLSTM has the lowest AUC (0.756) but the highest specificity, which is what a model with no external prior and only a 41-nt context should do—it picks up short sequential motifs that distinguish true sites from look-alikes.

Tower-removal confirms that none of the three is redundant. Pulling the RGCN out of the ensemble costs the most AUC (*−*0.68%), in line with its being the strongest single tower; pulling the BiLSTM costs the least (*−*0.27%), since its specificity-leaning bias is partly covered by RGCN. The range is narrow (0.27%–0.68%), which on its own could be read either as “all towers are useful” or as “the remaining two towers compensate.”

The CKA scores in the Discussion settle the question: the three representations are correlated but far from identical (CKA 0.55–0.67), so each tower carries information the other two do not, and the narrow AUC range is a sign of genuine complementarity rather than redundancy.

Fig 5 reports component-level ablations. Removing label smoothing produces the largest AUC decline (*−*1.06%), confirming its importance for calibrating predictions under severe class imbalance. Replacing BellPooling with mean pooling yields a minimal decline (*−*0.27%), indicating that the bell-shaped initialization captures most of the benefit with little learned deviation from the prior (Fig 9). Removing EMA slightly improves AUC (+0.15%), which is within noise range, indicating negligible impact on generalization under the current training regime.

**Fig 5.**
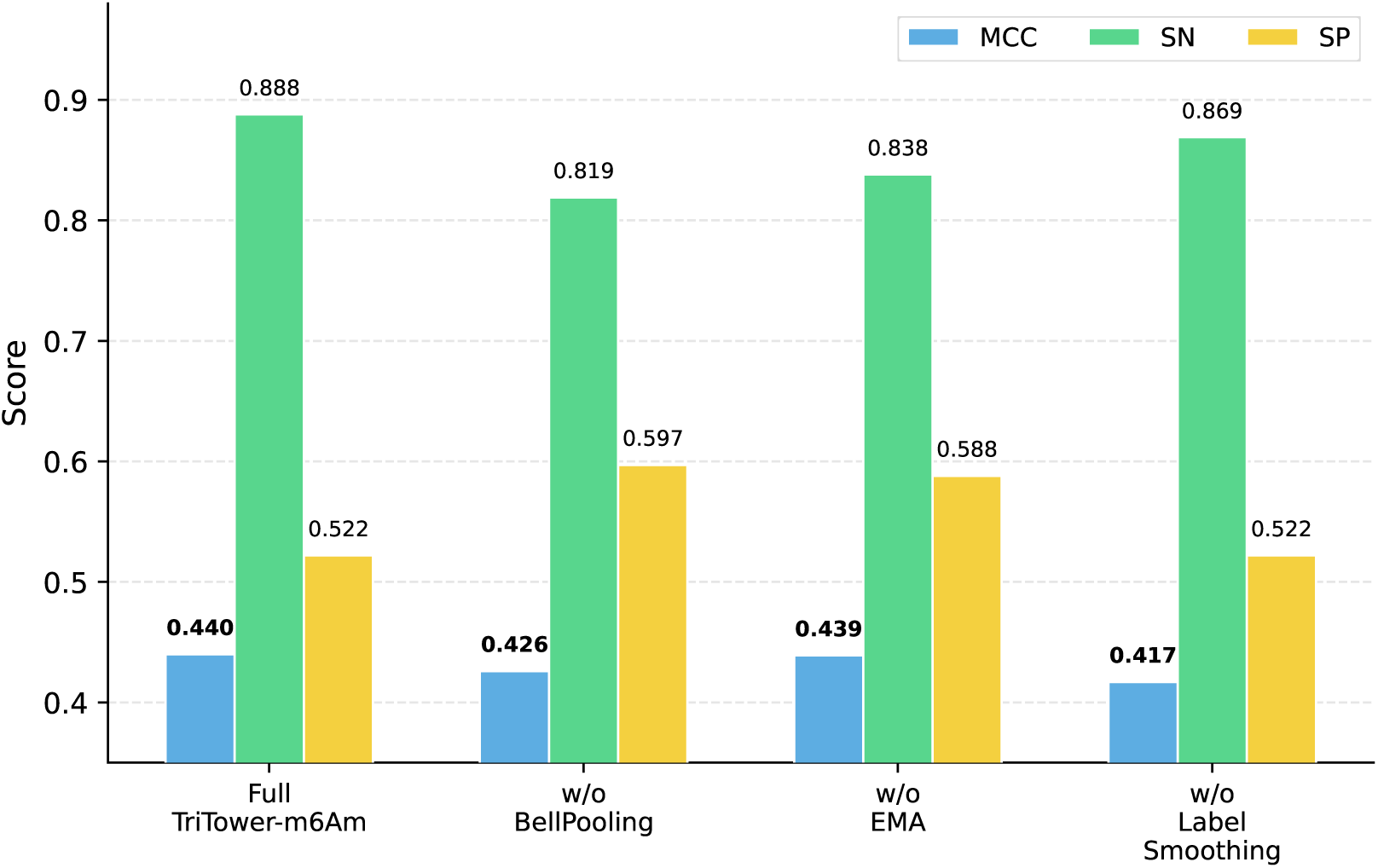
Component ablation study. “w/o” indicates the corresponding module is removed from the full model. AUC is omitted as it varies negligibly across configurations; MCC, SN, and SP reveal the distinct behavioral changes induced by each component.

A natural concern is whether the ensemble’s superior performance simply reflects increased model capacity rather than genuine complementarity among the three representations. To rule out this confound, we trained parameter-matched “wide” single-tower baselines in which each tower’s hidden dimension was enlarged to approximate the total parameter count of the full three-tower ensemble (*∼*330K). Specifically, the wide RNA-FM tower doubles the projection dimension from 96 to 160 and the multi-scale CNN channels from 48 to 80 per branch (yielding a merged width of 240 instead of 144), totaling 318K parameters; the wide BiLSTM tower doubles the hidden dimension from 64 to 128 per direction (256 total), totaling 549K parameters; and the wide RGCN tower triples the hidden dimension from 64 to 192, totaling 381K parameters. Each wide tower matches or exceeds the full ensemble in capacity (Fig 6).

**Fig 6.**
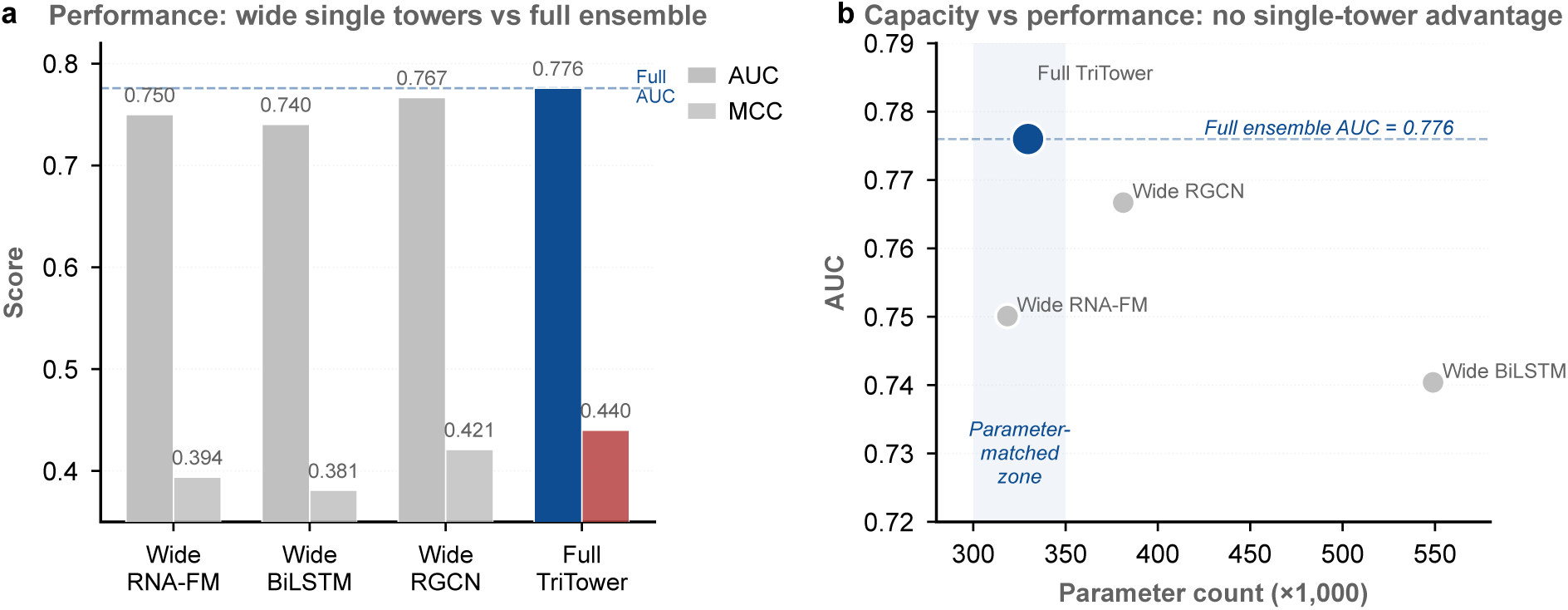
Parameter-matched single-tower baselines vs. full ensemble. (a) AUC and MCC on the independent test set. All three wide single towers fall below the full ensemble despite matching or exceeding its parameter count. (b) Capacity–performance scatter: increasing parameters alone does not recover ensemble performance. The shaded region marks the parameter-matched zone (*∼*300–350K).

On the independent test set, all three wide single towers underperform the full ensemble (AUC = 0.776, MCC = 0.440): wide RNA-FM reaches AUC = 0.750/MCC = 0.394, wide BiLSTM reaches AUC = 0.740/MCC = 0.381, and wide RGCN reaches AUC = 0.767/MCC = 0.421. The pattern is the same in every case: the wide tower matches or exceeds the full ensemble in parameter count, yet falls short on the test set. Two of the three (wide BiLSTM and wide RGCN) reach out-of-fold AUC above 0.90 on the training data and then drop to 0.74–0.77 on held-out data, which is the classical overfitting signature—extra capacity memorizes the training folds instead of learning a more general function. We read this as the clearest evidence in the paper that the ensemble’s gain is not a capacity effect: widening a single tower is strictly worse than splitting the same parameter budget across three heterogeneous representations, even though the latter option is shallower per representation.

### Structural mechanisms: edge type and node feature ablation

A key advantage of the RGCN’s relation-specific design is that it enables direct assessment of which structural relationships matter most. Table 6 reports two complementary sets of experiments: removing one edge type at a time from the full graph (columns 2–4), and using a single edge type in isolation (columns 5–7).

**Table 6.**
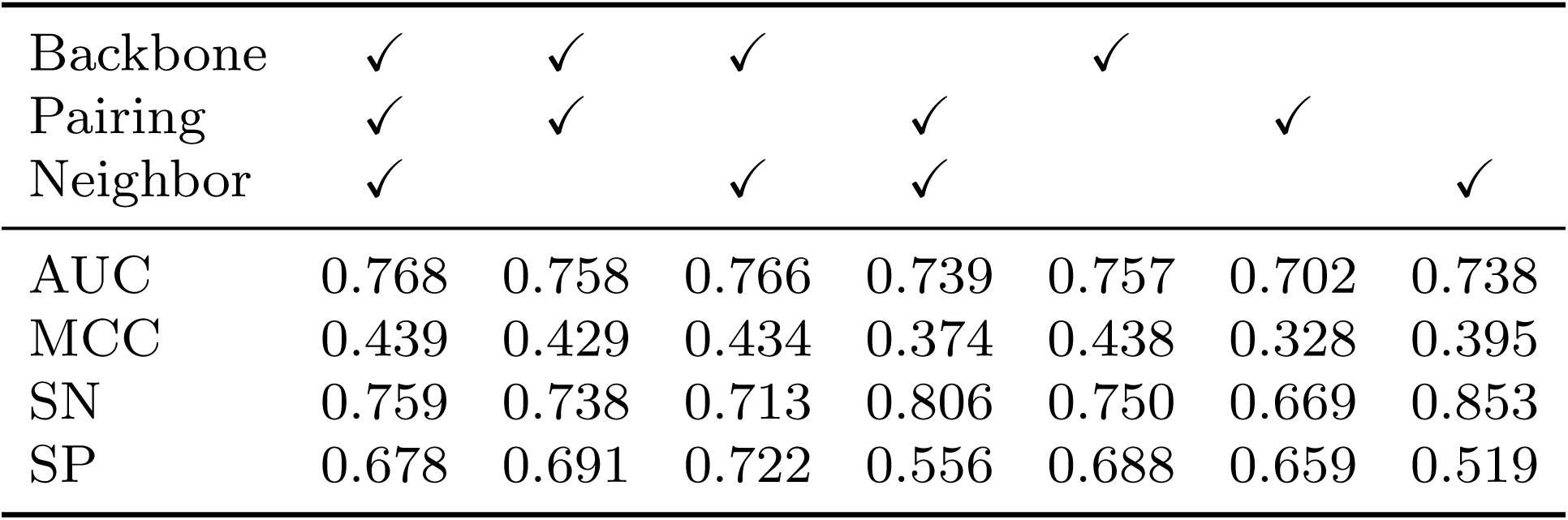
Edge type ablation for the RGCN tower. The first column lists the three edge types (backbone, pairing, neighbor); a checkmark indicates that the edge type is included in the corresponding model configuration.

**Fig 7.**
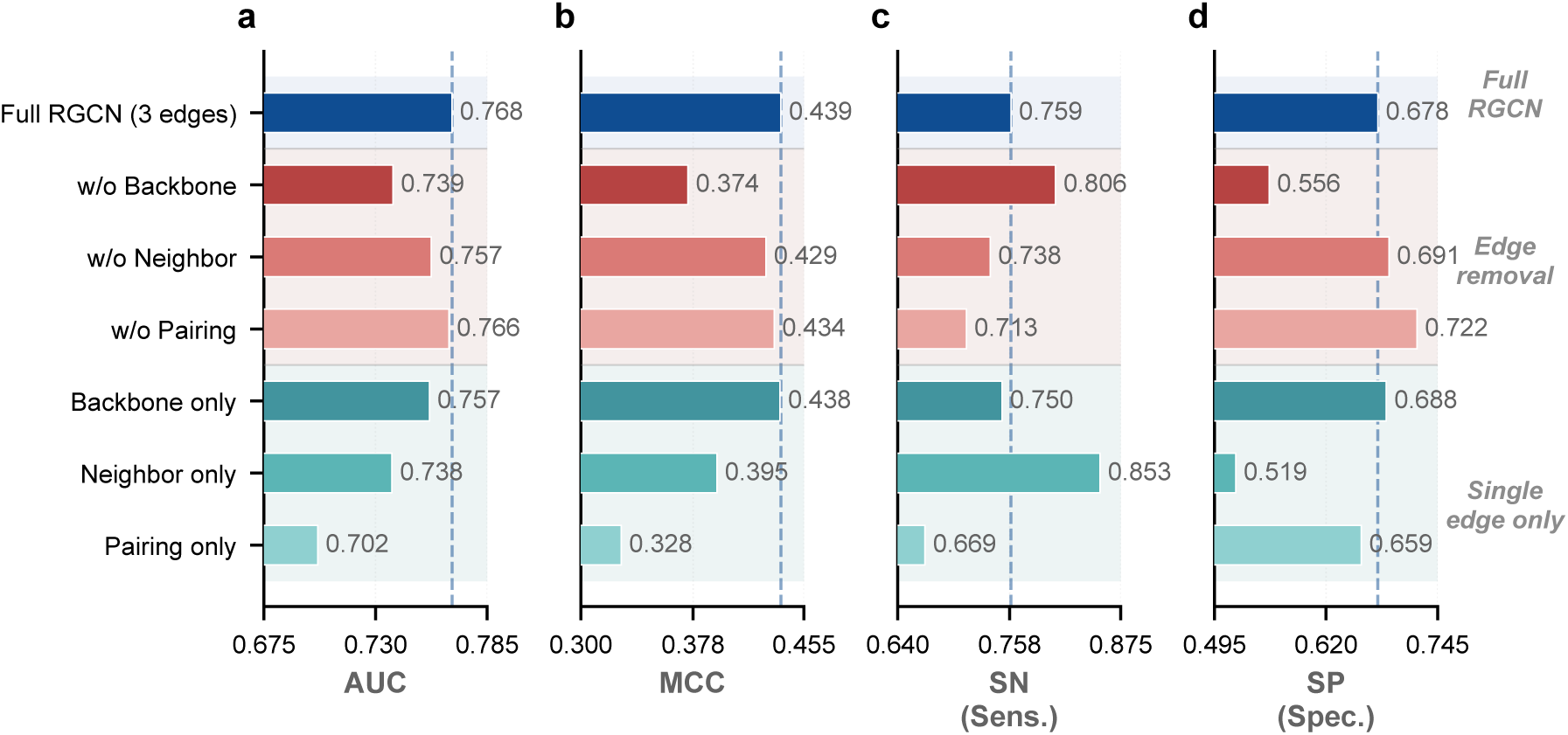
Edge-type ablation: per-metric comparison. Top group: full RGCN and removal of each edge type (red). Bottom group: single-edge-type-only models (teal). The dashed vertical line marks the full three-edge RGCN performance.

The structural signal is concentrated in the backbone. Pulling backbone edges out of the full graph costs 2.91% AUC (0.768 *→* 0.739) and 0.065 MCC, the largest single-edge drop in the table. Running the RGCN on backbone edges alone, with both pairing and neighbor edges deleted, still reaches AUC = 0.757 and MCC = 0.438—within 0.011 AUC and 0.001 MCC of the full three-edge model. In other words, almost everything the RGCN learns about m6Am can be learned from linear adjacency alone.

Pairing edges contribute the least. Dropping them from the full graph shifts AUC by only 0.002 (0.768 *→* 0.766), and a model that uses pairing edges in isolation is the weakest single-edge configuration (AUC = 0.701, MCC = 0.328). Neighbor edges sit in between: removing them costs 0.010 AUC, and neighbor-only reaches AUC = 0.738 with a sensitivity-leaning profile (SN = 0.853, SP = 0.519), i.e. it is willing to call positives but cannot reliably reject negatives.

Taken together, the three edge types form a clear hierarchy (backbone *≫* neighbor *>* pairing). This ordering is only visible because RGCN keeps edge types separate; a GAT with shared attention would average the three contributions and report a single structural effect, which is precisely the information loss that motivated the RGCN choice.

The edge-type ablation showed that pairing edges contributed negligibly to performance, which suggests that structural information might be carried by the node features rather than propagated through the edges. To test this, we performed a leave-one-group-out (LOGO) ablation on the 21-dimensional RGCN node feature vector, which comprises eight biologically motivated feature groups (Table 7). Each group was removed in turn while retaining the remaining seven groups, and the RGCN tower was retrained under identical settings (5-fold cross-validation, seed 123).

**Table 7.**
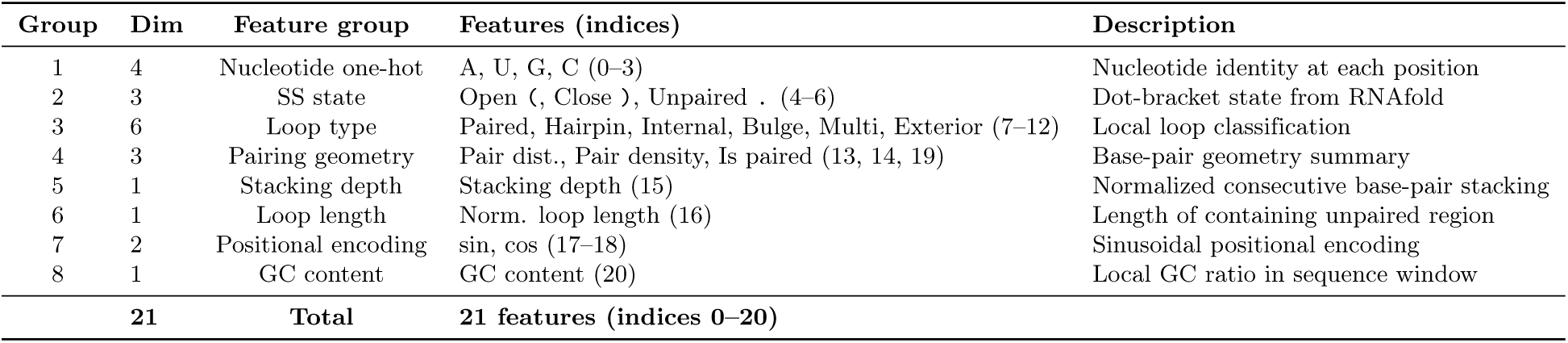
Composition of the eight LOGO ablation groups for the 21-dimensional RGCN node feature vector. Each group was removed in turn while retaining the remaining seven groups. Feature indices refer to columns in the 21-dimensional node feature vector defined in Table 2.

Fig 8 reports the results. The full 21-feature model achieved AUC = 0.764 and MCC = 0.438, consistent with the standalone RGCN tower performance reported in Table 6. Two feature groups dominated performance: removing nucleotide one-hot caused the largest drop (AUC: 0.764 *→* 0.698, ΔAUC = *−*0.067; MCC: 0.438 *→* 0.335, ΔMCC = *−*0.103), confirming that sequence identity remains the primary signal for m6Am recognition. Positional encoding was the second most important group (AUC: 0.764 *→* 0.724, ΔAUC = *−*0.040), reflecting the RGCN’s reliance on position information to locate the central modification site (position 20).

**Fig 8.**
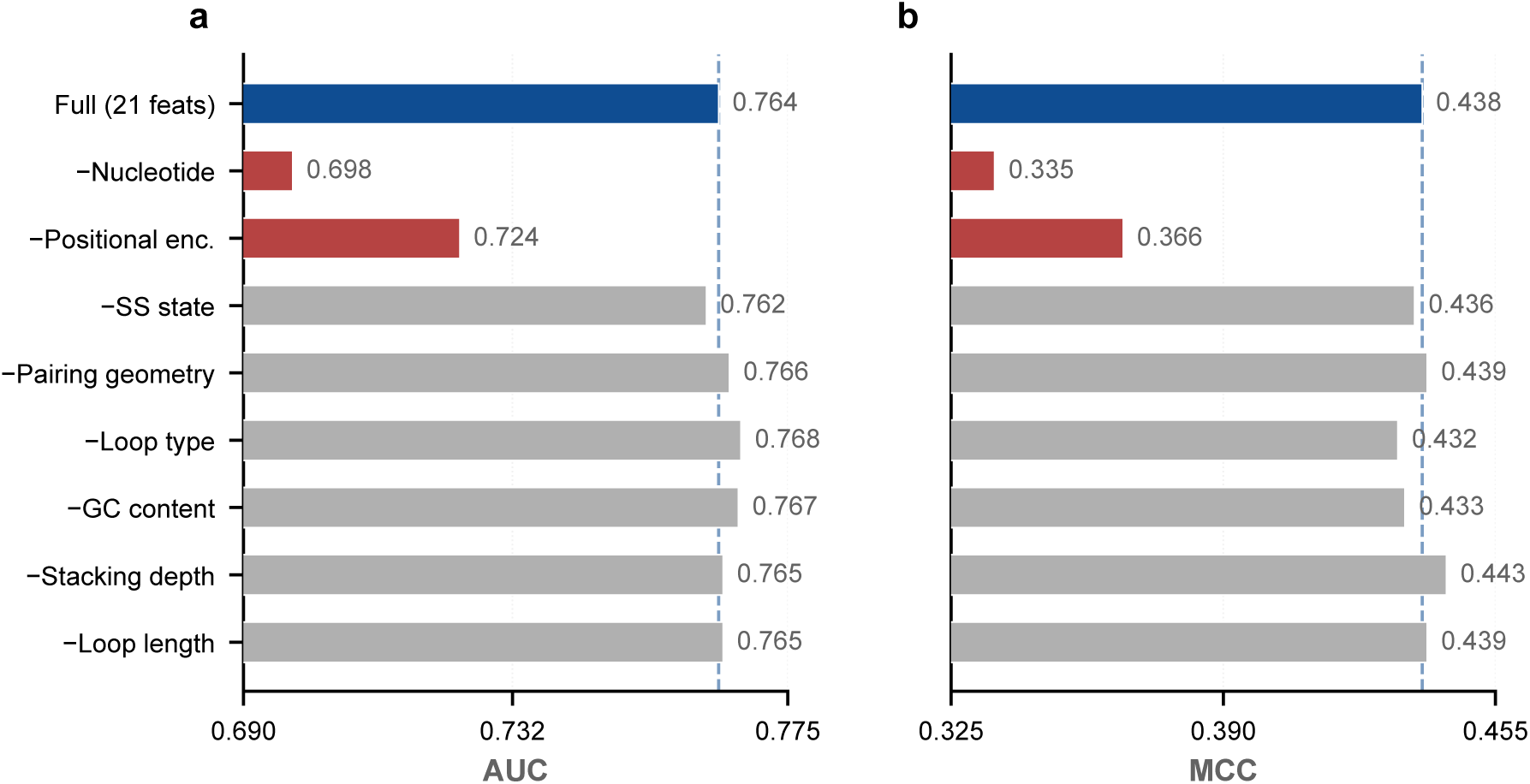
Node feature ablation for the RGCN tower. Leave-one-group-out ablation on the 21-dimensional node feature vector. The dashed line marks the full-model performance. Nucleotide one-hot and positional encoding are the two critical groups (red), while SS state and pairing geometry (the features redundant with pairing edges) show negligible impact (grey).

Removing the SS state or pairing geometry features caused negligible performance changes (SS state: ΔAUC = *−*0.002; pairing geometry: ΔAUC = +0.002), which resolves the puzzle raised by the edge-type ablation. Pairing edges contributed little not because base-pairing is irrelevant, but because the same information is already encoded in the node features: the dot-bracket state and loop-type vectors carry the same signal the pairing edges would propagate, so removing those edges deprives the model of information it can recover from the node attributes themselves. The remaining feature groups (loop type, GC content, stacking depth, loop length) likewise had minimal individual impact (*|*ΔAUC*| <* 0.004), indicating that the structural signal is distributed across multiple redundant encodings rather than concentrated in a single feature.

### Interpretability: BellPooling, IG attribution, and ensemble weights

The RNA-FM tower employs BellPooling to aggregate RNA-FM semantic features across the 41 positions into a single representation. Fig 9 compares the initialized and learned BellPooling weights. The initialized weights follow a symmetric bell curve centered at position 20 (the m6Am site), encoding the prior that positions near the modification site carry the most relevant semantic information. After training, the weights remain peaked near the center but develop a slight asymmetry, with positions 18–19 receiving marginally increased weight relative to the symmetric prior.

**Fig 9.**
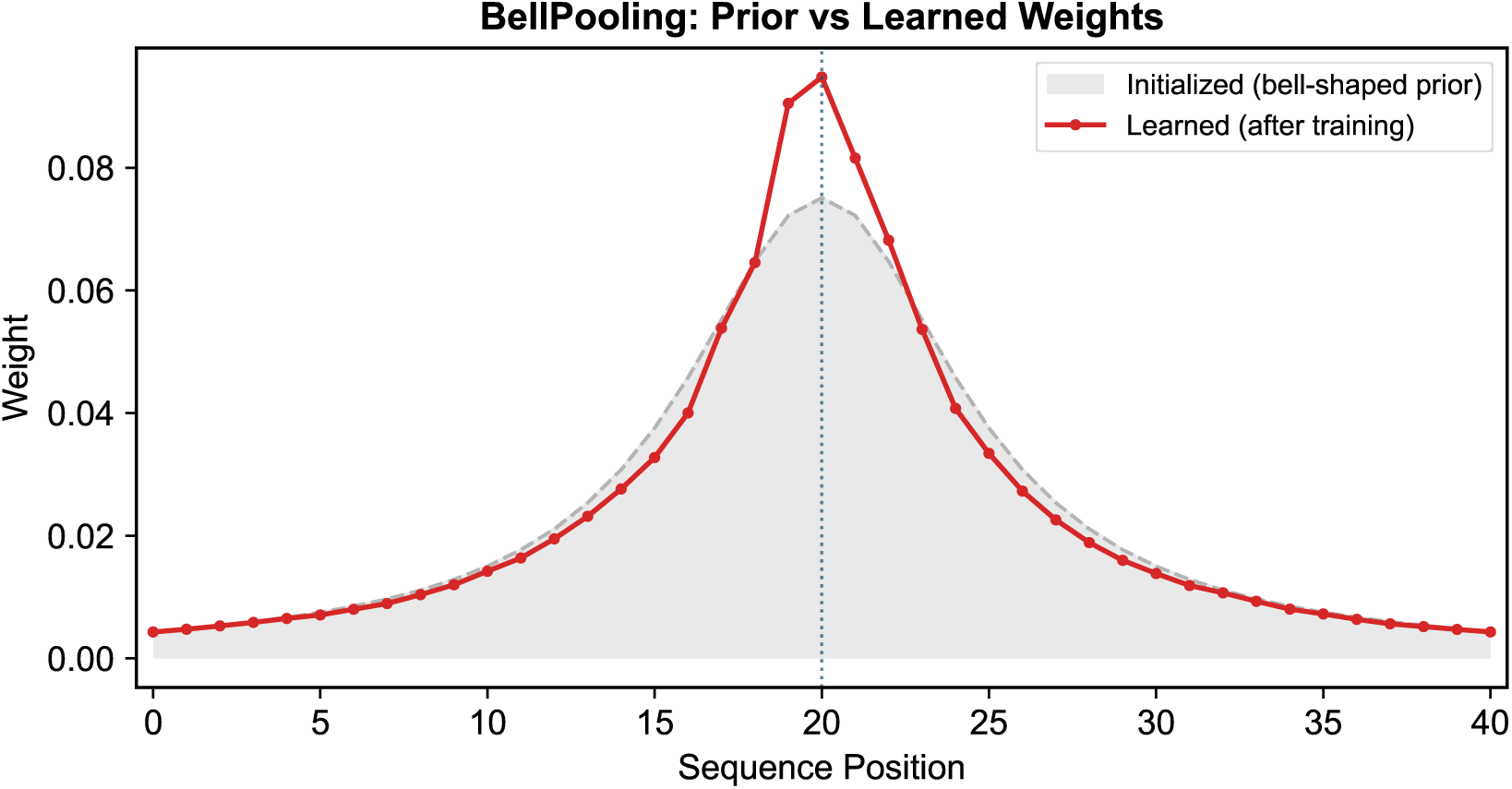
BellPooling weights: initialized (gray) vs. learned after training (red). The vertical dotted line marks position 20.

An important caveat: BellPooling operates on the 48-dimensional CNN-compressed RNA-FM features, not on the raw nucleotide sequence. The learned weights therefore reflect the model’s aggregation preference and should not be interpreted as direct sequence-level attribution. The stability of the center-peaked pattern after training indicates that the bell-shaped prior biases aggregation toward cap-proximal positions while permitting limited adaptation. In the ablation study (Fig 5), replacing BellPooling with uniform average pooling yielded a negligible AUC change (*−*0.27%), consistent with the observation that the learned weights deviated only slightly from initialization.

While BellPooling reveals the RNA-FM tower’s aggregation preference, it does not provide nucleotide-level attribution. To examine which positions the One-Hot BiLSTM tower relies on for its decisions, we applied Captum’s Integrated Gradients (IG) [30] (Captum v0.8.0) to the one-hot input of the BiLSTM tower. IG assigns each position an attribution score proportional to its contribution to the model’s output relative to a zero baseline (baselines=torch.zeros like(input), 50 integration steps). The sequence logos in Fig 10 were rendered with logomaker v0.8.7. Dropout layers in the BiLSTM were disabled during attribution to ensure deterministic gradients, while the model was kept in training mode to satisfy cuDNN’s LSTM backward-pass requirement.

**Fig 10.**
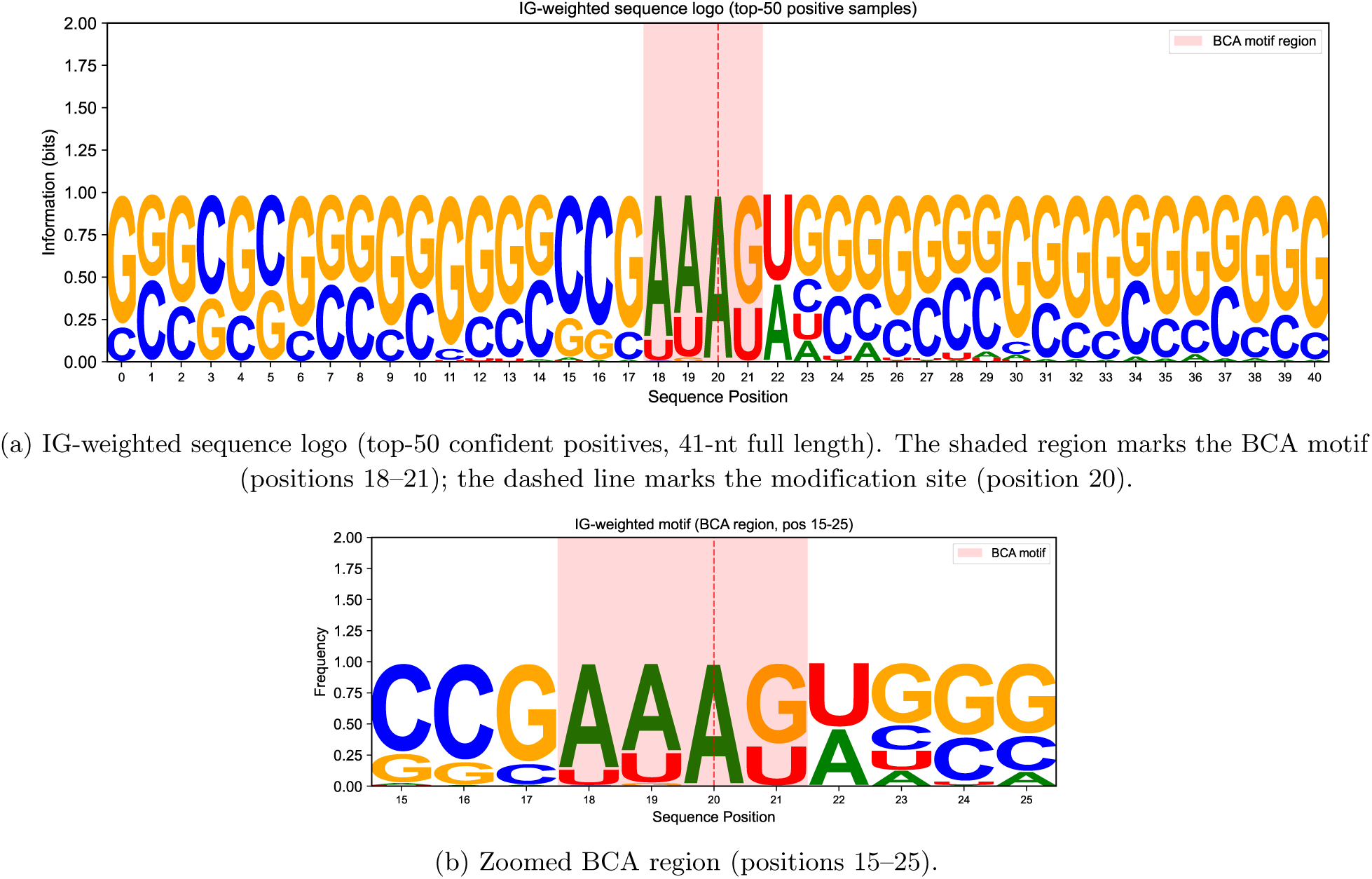
IG-weighted sequence logo from the One-Hot BiLSTM tower. Letter heights reflect IG-attribution-weighted nucleotide frequencies across the top-50 highest-confidence positive samples, not raw sequence frequencies. The peak attribution at position 22 indicates that the model’s discriminative signal lies downstream of the fixed BCA motif, consistent with PCIF1’s context-dependent methylation efficiency.

To extract a model-attribution-driven motif rather than a simple frequency logo, we selected the top-50 highest-confidence positive samples (ensemble probability *|*0.5*|* largest) and built a position weight matrix in which each position’s nucleotide contribution is weighted by its positive-part IG attribution. The resulting 41-nt logo (Fig 10) reveals two findings. First, the total IG weight concentrates downstream of the BCA motif: the peak attribution falls at position 22 rather than at the modification site itself (position 20), and the BCA region (positions 18–21) accounts for 21.3% of the total weight. Second, position 20 is fixed to adenosine by dataset construction in all samples, so the logo at this position is non-discriminative by design; the discriminative signal instead emerges in the flanking context, particularly the +2 downstream position. This pattern is biologically consistent with PCIF1’s substrate recognition: the enzyme binds the BCA motif [3] but its methylation efficiency is modulated by the local sequence context immediately downstream of the cap-proximal adenosine.

If the three towers encode genuinely complementary information, they should each contribute meaningfully: no single tower should dominate. AUC-weighted ensemble voting makes this directly testable, since each weight quantitatively measures a tower’s discriminative power on the training distribution.

The learned weights are RNA-FM (31.6%), One-Hot BiLSTM (34.5%), and RGCN (33.9%) (Fig 11). This near-uniform distribution indicates that all three towers carry independent predictive value; no single tower dominates the ensemble.

**Fig 11.**
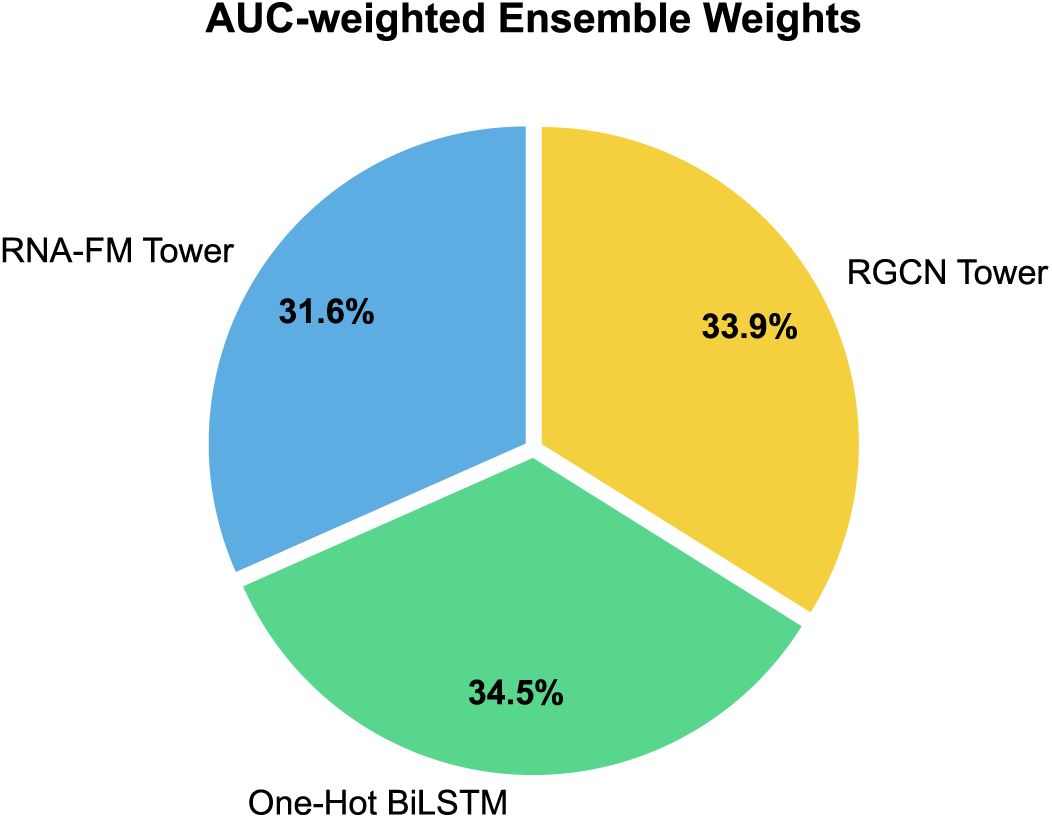
AUC-weighted ensemble weights for RNA-FM (31.6%), One-Hot BiLSTM (34.5%), and RGCN (33.9%).

Unlike black-box strategies such as stacking with a meta-learner, these weights provide an auditable decision trace. For any individual prediction, one can inspect how much each tower contributed to the final probability. This transparency is rare in deep learning ensembles and makes TriTower-m6Am more interpretable in practice.

### Robustness and generalization

To verify that TriTower-m6Am’s performance is not an artifact of a particular random seed, we retrained the full pipeline with three independent seeds (123, 42, 52) using identical hyperparameters and 5-fold cross-validation. The results (Table 8) demonstrate strong stability: AUC varies by only *±*0.003 (coefficient of variation = 0.34%) and MCC by *±*0.006 (CV = 1.46%). All three seeds outperform DTC-m6Am (AUC = 0.765, MCC = 0.411) in both metrics. The sensitivity and specificity also remain stable across seeds (SN: 0.865 *±* 0.019, SP: 0.539 *±* 0.023), confirming that the model’s decision behavior is robust to initialization randomness. Note that the MCC values in Table 8 differ slightly from the main result (Table 4, MCC = 0.440) because the multi-seed analysis reports metrics at a fixed optimal threshold determined per seed, whereas the main result uses the optimal threshold from the primary seed (123) for consistency with published benchmarks.

**Table 8.** Multi-seed stability analysis. Results from three independent seeds (123, 42, 52), each with 5-fold cross-validation.

| Seed | AUC | MCC | ACC | F1 | SN | SP |
| --- | --- | --- | --- | --- | --- | --- |
| 123 | 0.776 | 0.433 | 0.700 | 0.748 | 0.891 | 0.509 |
| 42 | 0.776 | 0.430 | 0.706 | 0.742 | 0.847 | 0.566 |
| 52 | 0.770 | 0.418 | 0.698 | 0.740 | 0.856 | 0.541 |
| Mean ± Std | **0.774 ± 0.003** | **0.427 ± 0.006** | **0.702 ± 0.003** | **0.743 ± 0.004** | **0.865 ± 0.019** | **0.539 ± 0.023** |
| CV (%) | 0.34 | 1.46 | 0.48 | 0.47 | 2.18 | 4.27 |

We first evaluated how TriTower-m6Am responds to varying training data availability by sub-sampling negatives at five ratios (1:1, 3:1, 5:1, 7:1, 10:1), keeping all 3,700 positives fixed. The independent test set (640 samples, balanced) remained unchanged. Results are reported in Table 9. Because sub-sampling negatives changes both the class balance and the total dataset size, this experiment measures sensitivity to *data availability* rather than class balance alone.

**Table 9.** Performance under varying training data availability. Higher ratios include more negative samples and larger total datasets.

| Ratio | $N_{\text{train}}$ | AUC | MCC | SN | SP | F1 | ACC |
| --- | --- | --- | --- | --- | --- | --- | --- |
| 1:1 | 7,400 | 0.745 | 0.394 | 0.831 | 0.547 | 0.728 | 0.689 |
| 3:1 | 14,800 | 0.743 | 0.388 | 0.800 | 0.578 | 0.720 | 0.689 |
| 5:1 | 22,200 | 0.769 | 0.416 | 0.738 | 0.678 | 0.716 | 0.708 |
| 7:1 | 29,600 | 0.759 | 0.403 | 0.719 | 0.684 | 0.707 | 0.702 |
| 10:1 | 40,700 | 0.776 | 0.428 | 0.828 | 0.588 | 0.739 | 0.708 |

Performance trends upward with dataset size, and the 10:1 ratio (the full training set) yields the best results (AUC = 0.776, MCC = 0.428). A modest dip occurs at the 7:1 ratio (AUC = 0.759, MCC = 0.403) relative to 5:1 (AUC = 0.769, MCC = 0.416), which may reflect subsampling variance. The 1:1 ratio performs similarly to 3:1 despite better class balance, indicating that the smaller negative pool (3,700 vs. 11,100 negatives) limits performance more than the balance advantage helps.

To isolate the effect of class reweighting from dataset size, we performed a controlled experiment in which all 40,700 training samples were used for every condition and only the positive class weight (pos weight) in BCEWithLogitsLoss was varied (Fig 12).

**Fig 12.**
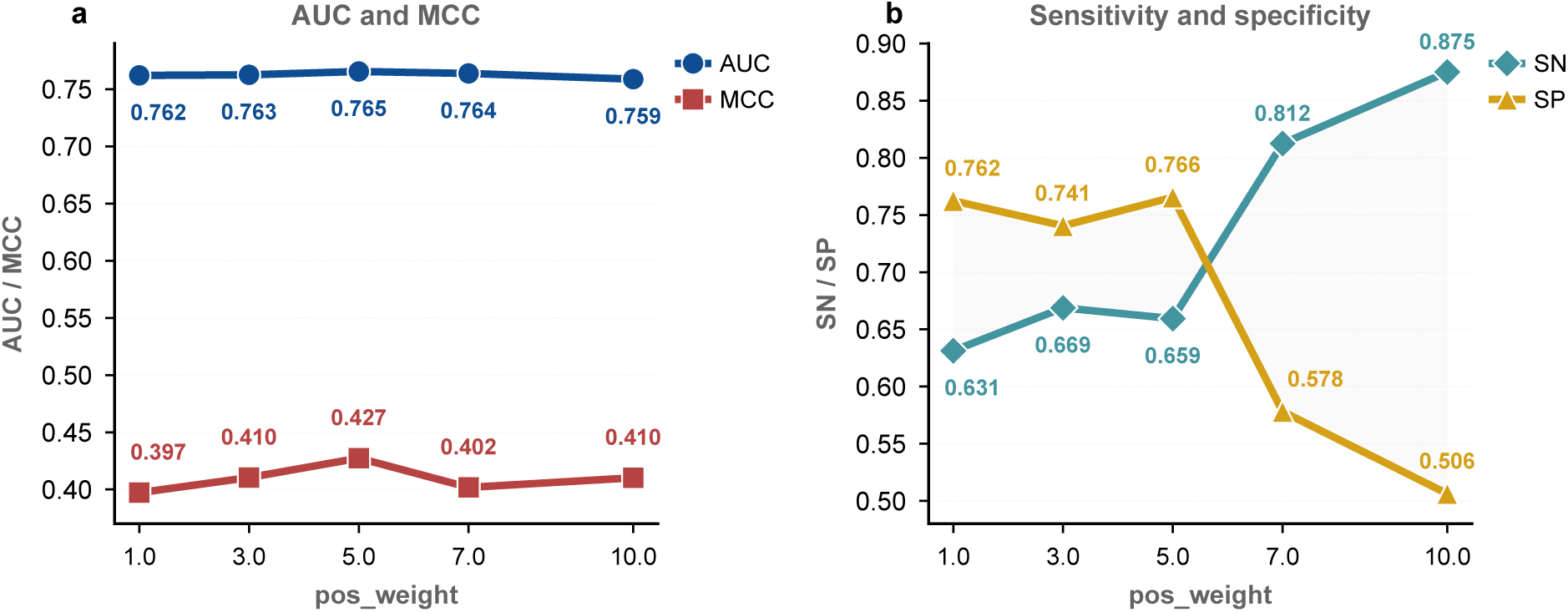
pos weight sensitivity with fixed dataset size. All 40,700 training samples used; only pos weight varies. (a) AUC and MCC. (b) Sensitivity and specificity.

Under fixed dataset size (all 40,700 training samples), AUC was largely insensitive to pos weight. Across the five tested values (1.0, 3.0, 5.0, 7.0, 10.0), AUC ranged from 0.759 to 0.766 (Δ = 0.007) and peaked at pos weight=5.0. MCC also peaked at pos weight=5.0 (0.427). Sensitivity and specificity, in contrast, showed a monotonic trade-off: sensitivity rose from 0.631 to 0.875 while specificity fell from 0.763 to 0.506 as pos weight increased from 1.0 to 10.0. At pos weight=10.0, specificity and AUC both reached their lowest values (0.506 and 0.759). The default cap of 3.0 (AUC = 0.763, MCC = 0.410, SN = 0.669, SP = 0.741) placed the operating point close to the AUC-optimal value of 5.0. The narrow AUC range across a 10*×* change in pos weight (Δ = 0.007) contrasts with the large shifts in sensitivity and specificity, indicating that the reweighting strategy primarily tunes the sensitivity–specificity balance rather than overall discriminative power.

Taken together, the two experiments show that (i) TriTower-m6Am benefits primarily from larger and more diverse training sets rather than from a specific class-balance configuration, and (ii) the default pos weight cap of 3.0 provides a near-optimal operating point: AUC and MCC peak at pos weight=5.0, but the difference from 3.0 is negligible (ΔAUC *<* 0.003). The model is robust to dataset-specific variation in negative-to-positive ratios, making it practical for real-world use across tissues, sequencing protocols, and annotation standards.

The balanced 1:1 test set (320 positive, 320 negative) does not reflect the extreme rarity of m6Am sites in genome-wide screening, where the ratio of true m6Am sites to candidate adenosine positions can reach 1:100 or higher. To evaluate model performance under realistic deployment conditions, we constructed an independent set of negative samples from the GENCODE v44 human reference transcriptome.

### External negative construction

We scanned all GENCODE v44 transcripts for the BCA motif (B = C/G/U), the canonical cap-proximal sequence context recognized by PCIF1 [3]. After removing known m6Am positive sites and deduplication, 8.6 million candidate BCA positions were identified; 50,000 were randomly sampled for evaluation. Each position was extracted as a 41-nt window (*±*20 nt flanking), and RNAfold was used to predict secondary structure. These BCA-derived negatives are entirely independent of both the training and test sets used in the main experiments. Note that the sampling is not stratified by transcript expression level or transcript length: because BCA motifs are scanned across the full transcript body, longer transcripts contribute proportionally more candidates, and no expression-weighting is applied. This sampling design matches the deployment scenario of a transcriptome-wide screen, where every candidate adenosine must be evaluated regardless of the source transcript’s abundance.

### Evaluation protocol

We combined the 320 original test positives with varying numbers of BCA negatives to construct three test sets at imbalance ratios of 1:1 (320 pos + 320 neg), 1:10 (320 pos + 3,200 neg), and 1:100 (320 pos + 32,000 neg). The trained 5-fold ensemble was applied without any retraining or threshold adjustment. We report AUC, average precision (AP), MCC, F1, sensitivity (SN), and specificity (SP) (Fig 13).

**Fig 13.**
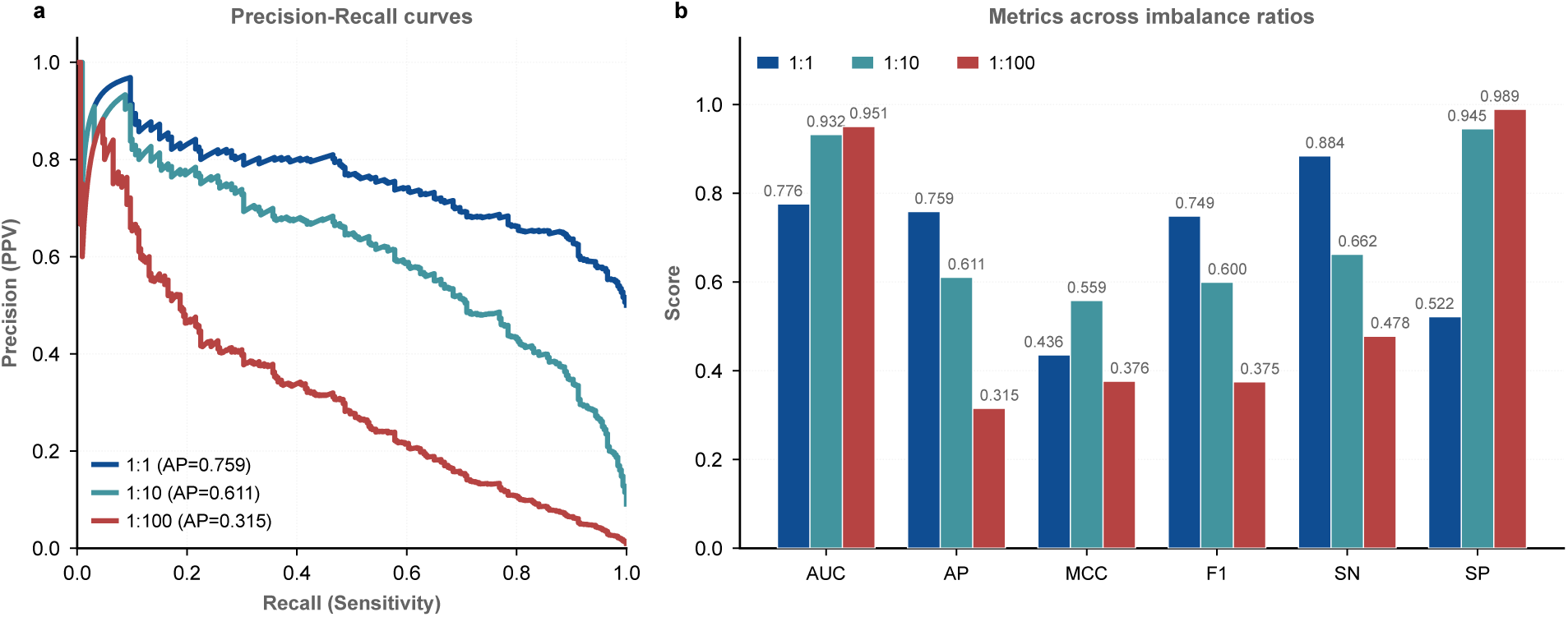
Performance under varying class imbalance ratios. (a) Precision-recall curves at 1:1, 1:10, and 1:100 ratios using external BCA-derived negatives from the GENCODE transcriptome. (b) Metrics comparison across the three ratios. AUC and SP increase with imbalance due to the inclusion of easier BCA negatives, while AP and SN decrease as the positive class becomes rarer.

## Results

AUC increased from 0.776 (1:1) to 0.933 (1:10) to 0.951 (1:100), reflecting the inclusion of BCA-derived negatives that the model can distinguish more easily than the original test negatives. In contrast, average precision decreased sharply from 0.759 (1:1) to 0.611 (1:10) to 0.315 (1:100), indicating that in genome-wide screening scenarios, the model produces a substantial number of false positives relative to true positives. Sensitivity decreased from 0.884 to 0.663 to 0.478 as the MCC-optimal threshold shifted upward (0.48 *→* 0.67 *→* 0.81), while specificity increased from 0.522 to 0.945 to 0.989. MCC peaked at the 1:10 ratio (0.559) before declining at 1:100 (0.377).

### Interpretation

The divergence between AUC and AP under extreme imbalance is expected and informative. AUC, which aggregates over all thresholds, benefits from the model’s ability to rank the easier BCA negatives below true positives. AP, which focuses on the precision of the top-ranked predictions, reveals the practical challenge: at 1:100, even a specificity of 0.989 yields 342 false positives among 32,000 negatives, exceeding the 320 true positives. This confirms that genome-wide m6Am screening requires complementary filtering (e.g., cap-proximal annotation, transcript start site databases) to reduce the candidate space before model-based prediction. The trained model generalizes to external data without retraining, and the MCC-optimal threshold adapts to the operating ratio, but users should adjust the decision threshold based on the target prevalence in their application. Practical mitigations for the AP drop under extreme imbalance include hard-negative mining to retrain on the highest-scoring false positives, cost-sensitive learning with explicit false-positive penalties, and posterior threshold calibration against an estimate of the operating-ratio prevalence. These directions are complementary to the cap-proximal pre-filtering noted above and are left for future work.

### Case studies: high-confidence predictions

To illustrate the ensemble’s behavior in practice, we examined the 10 most confident positive predictions (highest ensemble probability) and the 10 most confident negative predictions (lowest ensemble probability) on the independent test set (Figs 14 and 15).

**Fig 14.**
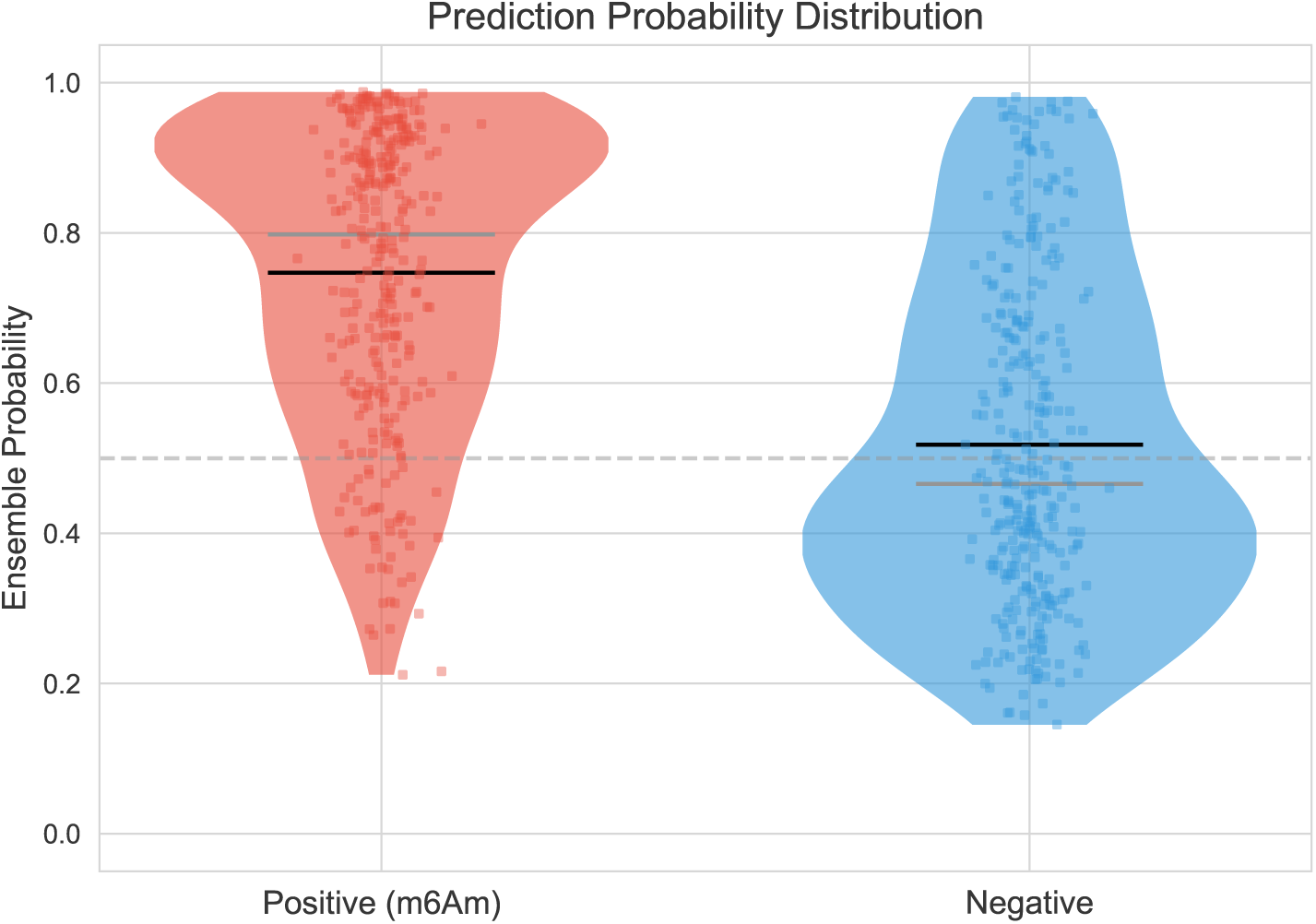
Distribution of ensemble probabilities for top-10 positive and top-10 negative predictions.

**Fig 15.**
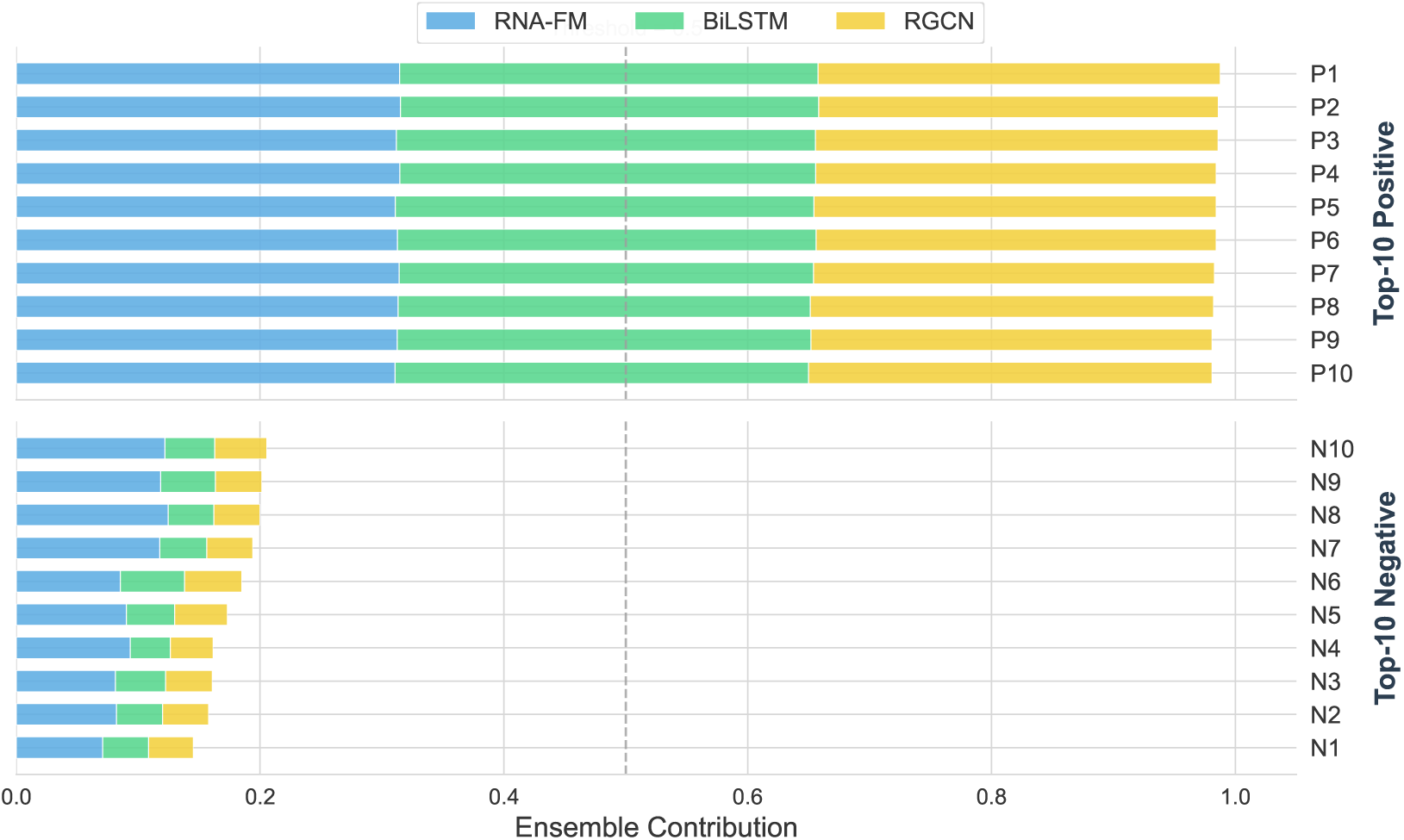
Tower contributions for top-10 positive and top-10 negative predictions.

High-confidence predictions require consensus across towers, since each tower captures a distinct signal. The top-10 positive predictions illustrate this: all three towers agree with high individual probabilities (RNA-FM: 0.98–0.99, One-Hot BiLSTM: 0.98–0.99, RGCN: 0.97–0.98), yielding a precision of 90% (9 true positives, 1 false positive). No single tower dominates these predictions.

Among the top-10 negative predictions, all 10 are true negatives (ensemble probabilities 0.15–0.21). Here the towers exhibit an asymmetric confidence pattern: the One-Hot BiLSTM and RGCN towers are highly confident (probabilities 0.09–0.15), while the RNA-FM tower is markedly less certain (probabilities 0.22–0.39). Despite this asymmetry, the AUC-weighted ensemble correctly classifies all 10 samples as negative, indicating that the voting scheme is robust to individual tower uncertainty: the two confident towers override the RNA-FM tower’s relatively higher probabilities, preventing false positives in the high-confidence negative regime.

## Discussion

### From single representation to representation complementarity

The complementarity among semantic, sequential, and structural representations explains the performance gain over single-representation predictors. We did not assume these channels were complementary; we designed the architecture so the question could be tested. Pairwise CKA analysis gives a partial answer: RNA-FM vs. One-Hot BiLSTM = 0.547, RNA-FM vs. RGCN = 0.655, and One-Hot BiLSTM vs. RGCN = 0.668 (Fig 16). Read in isolation, a CKA near 0.5–0.7 might look like moderate overlap, but in the context of the tower-removal ablation the number is informative rather than reassuring: pulling any single tower out of the ensemble costs AUC, and the CKA scores are high enough that the remaining towers compensate partially but low enough that they cannot compensate fully. The ensemble gain is therefore genuine complementarity, not a capacity artifact—a reading that the wide single-tower baselines confirm independently (see the wide-baseline analysis above). The BellPooling mechanism, which applies a center-focused positional prior to the RNA-FM tower output, was retained for its conceptual simplicity despite offering negligible performance benefit over uniform pooling (Fig 5); the learned weights deviate only slightly from the bell prior, which we read as the model confirming rather than overriding the biological expectation that cap-proximal positions matter most.

**Fig 16.**
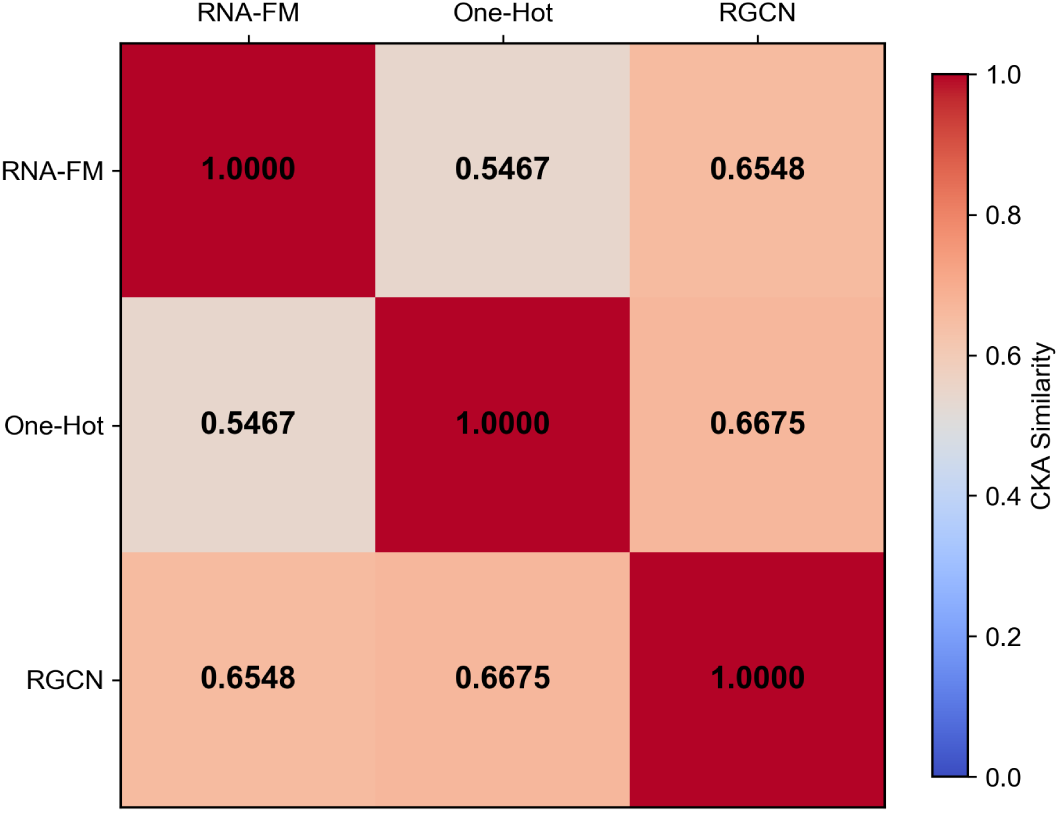
Pairwise CKA similarity between tower representations.

### Structural signal is dominated by backbone adjacency

The RGCN edge-type ablation reveals that backbone adjacency, rather than long-range pairing, dominates the structural signal in m6Am prediction. We replaced the Graph Attention Network (GAT) used in preliminary experiments with a Relational Graph Convolutional Network (RGCN). GAT assigns type-agnostic attention shared across all edge types. RGCN, in contrast, uses relation-specific weight matrices, enabling differentiated message passing for backbone, base-pairing, and neighbor edges. The resulting edge-type hierarchy (backbone *≫* neighbor *>* pairing) would be invisible to GAT, which is the concrete reason the replacement matters: without separate edge types, the small but real contribution of pairing and neighbor edges would be averaged into a single “structural” number, and the design lesson—that linear adjacency already carries most of the structural signal—would be lost.

The edge ablation turns up a result we did not anticipate. Backbone edges alone reach AUC = 0.757 and MCC = 0.438, within 0.011 AUC of the full three-edge model (AUC = 0.768, MCC = 0.439); pairing edges alone reach only AUC = 0.701. We expected pairing edges to matter, on the reasoning that PCIF1 recognizes a structured substrate, but the data point the other way: PCIF1 recognizes the cap-proximal adenosine in its local linear context [2, 3], and the backbone carries enough of that context for the RGCN to work. The minimal contribution of base-pairing edges is attributable to two factors. First, secondary structure information is already encoded in the node features (dot-bracket notation and loop type), so the pairing edges carry signal the model can recover from the node attributes themselves. Second, m6Am is installed co-transcriptionally, before the nascent RNA has fully folded, which weakens any 2D-topology signal at the moment of recognition.

This raises an apparent paradox: the RGCN is the strongest single tower, yet the individual structural feature groups each contribute marginally. The node feature ablation (Fig 8) resolves it. The structural signal is not concentrated in any single feature group; it is distributed across SS state, loop type, pairing geometry, and the pairing edges, all of which carry overlapping information. Removing any one group therefore has minimal impact because the others compensate, but the cumulative effect of structural modeling still yields a stable advantage over non-structural representations. We read this distributed-redundancy pattern as the model coping with the fact that RNA secondary structure is an emergent property of the sequence: encoding it through several correlated feature groups makes the tower robust to noise in any single structure predictor output, which is a useful property given that RNAfold itself is only *∼*73% accurate per base pair.

### Built-in interpretability via AUC-weighted fusion

The AUC-weighted ensemble provides an inherently interpretable fusion, as each prediction can be audited by inspecting tower weights rather than treating the model as a black box. Interpretability is a property of the architecture, not a post-hoc addition. The typed RGCN edges let us ask which structural relationship a prediction rests on (the edge-type hierarchy in the ablation analysis above); the AUC-weighted ensemble weights let us ask how much each tower contributed to a given prediction (Fig 11). The two views answer different questions—one about graph structure, one about tower contribution—and neither requires a separate post-hoc explainer trained to approximate the model. In practice this means a researcher who disagrees with a prediction can look at the tower weights and the edge-type ablation and form a specific hypothesis about what the model is “seeing,” rather than re-running the entire pipeline with a different explainer. The cost is that the weights and edge types are coarse-grained: they do not reach nucleotide resolution, which is why we added the IG-weighted logo in the interpretability analysis above as a complementary, model-specific attribution.

### Limitations and future work

Several limitations of TriTower-m6Am warrant discussion.

First, the high sensitivity (0.888) comes at the cost of moderate specificity (0.522). In applications where false positives are costly (such as experimental validation pipelines), threshold adjustment or cost-sensitive learning may be preferable.

Second, the RGCN tower relies on computationally predicted secondary structures from RNAfold. These predictions have an estimated accuracy of *∼*73% for individual base pairs. Integrating experimental structure data from techniques such as SHAPE-MaP or DMS-seq could improve the reliability of structural features.

Third, the node feature ablation (Fig 8) revealed redundancy between the SS state/pairing geometry features and the pairing edges, suggesting that a more compact node representation could be designed in future work. We retained the full feature set to preserve the interpretability of the contribution analysis and avoid introducing changes that would require re-validating the entire pipeline.

Fourth, RNA-FM embeddings are extracted from a frozen model. End-to-end fine-tuning of the RNA-FM backbone was not explored due to GPU memory constraints: the base model requires approximately 16 GB for inference alone. Parameter-efficient methods such as LoRA could enable task-specific adaptation without exceeding memory limits.

Fifth, the model is trained and evaluated exclusively on human m6Am data. Although m6Am is conserved across vertebrates and PCIF1 is highly conserved across mammals, cross-species evaluation was not performed in this study for two reasons. First, experimentally validated m6Am site catalogs with single-nucleotide resolution remain scarce for non-human organisms, limiting the availability of independent test sets. Second, cross-species performance differences are difficult to interpret: a drop in performance could reflect genuine species-specific biology, but it could equally reflect biases in the reference transcriptome, differences in sequencing protocol, or systematic differences in 5’UTR length and GC content across species. To avoid confounding these factors, we prioritized the external BCA-derived negative evaluation (see the external evaluation above), which tests generalization to independent sequences within the same species while controlling for cap-proximal context. Dedicated cross-species evaluation with matched experimental protocols is an important direction for future work.

Sixth, the external generalization evaluation (see the external evaluation above) extends only the negative-sample distribution to the GENCODE transcriptome, while the positive sites still originate from the same batch of miCLIP experiments used in the main test set. Cross-platform and cross-laboratory generalization (covering independent experimental techniques, distinct cell lines, and alternative sequencing protocols) is therefore not yet verified. Conclusions about deployment-ready performance should be tempered accordingly: an independent positive-site catalog generated by a different miCLIP or m6Am-seq protocol is needed to confirm that the model generalizes beyond a single experimental source.

Future directions include incorporating additional modalities such as genomic context and evolutionary conservation scores. Exploring end-to-end fine-tuning of RNA-FM with parameter-efficient methods and developing multi-task learning frameworks for joint prediction of m6A, m6Am, and other RNA modifications could further enhance predictive power. Extending the triple-tower architecture to other RNA modification prediction tasks (e.g., m5C [31], Ψ [32], m1A [33]) will test its generalizability [34], and conducting experimental validation of high-confidence novel predictions will assess real-world discovery utility.

## Conclusion

We presented TriTower-m6Am, a triple-tower architecture that combines three representations of an RNA window—semantic (RNA-FM), sequential (One-Hot BiLSTM), and structural (RGCN with typed edges)—for m6Am site prediction. On the independent test set the model reaches AUC = 0.776, MCC = 0.440, and SN = 0.888, improving on DTC-m6Am most visibly in sensitivity. The 8.8 percentage-point gain in SN is the number we would foreground: in a screening pipeline each missed m6Am site is a separate wet-lab experiment, so catching roughly 9 additional sites per 100 real ones is the practical return of the architecture.

The complementarity of the three towers is supported on two independent grounds. The CKA scores (0.55–0.67) are high enough that no tower is redundant but low enough that none is dispensable, and the parameter-matched wide single-tower baselines fall below the ensemble despite matching or exceeding its capacity—so the gain is not a capacity effect but a representation effect. The AUC-weighted voting weights are readable from the architecture, which makes each tower’s contribution to a prediction auditable without a post-hoc explainer.

Two findings from the ablations changed how we think about the problem. The RGCN’s typed edges expose a structural hierarchy in which linear backbone connectivity carries almost all of the structural signal, while base-pairing edges contribute little—a result that lines up with the co-transcriptional timing of PCIF1-mediated methylation and that would be invisible to a model with shared, type-agnostic attention. And the node feature ablation shows that the structural signal is distributed across several redundant encodings rather than concentrated in one, which we read as the model hedging against the noise of a single structure predictor. Extending the triple-tower pattern to other RNA modifications would test whether this distributed-redundancy property generalizes, or whether it is specific to cap-proximal recognition.

## List of abbreviations

m6Am: N^6^,2’-O-dimethyladenosine
m6A: N^6^-methyladenosine
m7G: 7-methylguanosine
mRNA: messenger RNA
PCIF1: Phosphorylated CTD Interacting Factor 1
CAPAM: Cap-Specific Adenosine Methyltransferase
FTO: Fat mass and obesity-associated protein
miCLIP: methylation individual-nucleotide-resolution cross-linking and immunoprecipitation
RNA-FM: RNA Foundation Model
BiLSTM: Bidirectional Long Short-Term Memory
RGCN: Relational Graph Convolutional Network
GAT: Graph Attention Network
AUC: Area Under the ROC Curve
ROC: Receiver Operating Characteristic
PR: Precision-Recall
AP: Average Precision
MCC: Matthews Correlation Coefficient
SN: Sensitivity
SP: Specificity
F1: F1 Score
ACC: Accuracy
EMA: Exponential Moving Average
OOF: Out-of-Fold
AMP: Automatic Mixed Precision
CNN: Convolutional Neural Network
CKA: Centered Kernel Alignment
GELU: Gaussian Error Linear Unit
LoRA: Low-Rank Adaptation.

## Acknowledgments

Not applicable.

## Declarations

### Ethics approval and consent to participate

Not applicable.

### Consent for publication

Not applicable.

### Availability of data and materials

The source code and pre-trained models are available at https://github.com/xiong-0212/TriTower-m6Am. The benchmark dataset is derived from Huang et al. [10].

### Authors’ contributions

K.X. performed the experiments, analyzed the data, and drafted the manuscript. J.J. conceived and supervised the study, revised the manuscript, and approved the final version.

## Supporting information

**S1 Table.** Computational efficiency of each tower and the ensemble. Inference time and peak GPU memory are averaged over 10 runs on a single NVIDIA GPU (16 GB) with batch size 1. Parameters are the trainable parameter count of each tower’s downstream encoder (RNA-FM backbone is frozen and excluded).

| Component | Parameters | Inference time (ms) | Peak GPU memory (MB) |
| --- | --- | --- | --- |
| RNA-FM tower | 140,974 | 0.65 | 10.63 |
| One-Hot tower | 143,489 | 0.47 | 42.35 |
| RGCN tower | 45,203 | 4.38 | 10.53 |
| TriTower ensemble | 329,666 | 6.18 | 42.35 |

